# Mycobacterial infection uncovers plasticity of Kupffer cells

**DOI:** 10.1101/2025.02.07.636999

**Authors:** Jana Neuber, Florens Lohrmann, Samuel Wald, Merve Göçer, Anne Kathrin Lösslein, David Obwegs, Vitka Gres, Torsten Goldmann, Manuel Rogg, Christoph Schell, Sebastian Preißl, Sagar, Philipp Henneke

## Abstract

*Bona fide* Kupffer cells (KCs) are prenatally seeded and show unique functional and immunophenotypic features among tissue macrophages. They are considered as terminally differentiated, and adaptability in disease is attributed to recruited, monocyte-derived KCs. Here, we investigated the extent of KC plasticity and the impact of origin in mycobacterial infections that target macrophages and can persist for months. Fate-mapping combined with high-resolution imaging revealed the emergence of a unique, infection specific KC subset which downregulated the signature markers CLEC4F and VSIG4 (“KC^low^”). KC^low^ were derived from *bona fide* KCs and located exclusively to granuloma cores. In contrast, monocyte-derived macrophages were contained at the granuloma borders and contributed to this tissue reaction. ATAC and single-cell RNA sequencing identified a specific signature of KC^low^ with high antimycobacterial activity and specialization to a hypoxic microenvironment. Despite their fundamental deviation from the classical KC phenotype, KC^low^ showed remarkable adaptability, and were capable to return to a homeostatic-like KC state. Accordingly, mycobacterial infections unmask KCs as highly plastic cells, capable of responding to extreme environmental changes.

## Main

Tissue-resident liver macrophages, called Kupffer cells (KCs), are prenatally seeded by either progenitors from the yolk sac or fetal liver^1^. Due to their distinct anatomical localization within the sinusoids, they are poised to capture blood-borne pathogens^2^. In addition, they play pivotal roles in the clearance of senescent erythrocytes, as well as gut-derived microbial products^3^.

In homeostasis, KCs self-renew and receive negligible input from monocytes^1^. However, during liver damage, infection-mediated cell death or after experimental KC depletion, monocytes enter the liver in high numbers and differentiate into monocyte-derived KCs (MoKCs) with the acquisition of a KC specific marker set including CLEC4F, VSIG4, or TIM-4^4,5^. The upregulation dynamics of KC markers in MoKCs vary, e.g. expression of CLEC4F can be detected after around one week, while TIM-4 is only partially expressed after one month in experimental KC depletion^6^.

The initiation of KC differentiation requires KCs to extend protrusions into the space of Disse, where they interact with liver sinusoidal endothelial cells (LSECs) and hepatic stellate cells (HSCs)^4,7,8^. Accordingly, KCs are prime examples of terminally differentiated macrophages with highest specification to their tissue niche early in life. In line with this morphological uniqueness, they are the only macrophages that express CLEC4F^6^.

Like other tissue macrophages, KCs are targeted by mycobacteria, which chronically perturb local innate immunity^9,10^. Mycobacteria have specifically adapted to macrophages in a coevolution extending over tens of thousands of years^11^. Thus, they have developed mechanisms to not only persist in this, in principle, hostile environment^12,13^, but to use macrophages as a proliferative niche^14^. Infections with mycobacteria lead to the formation of granulomas, highly dynamic cellular structures consisting of specialized macrophage subsets, in particular multinucleated giant cells (MGCs), foamy macrophages, or epithelioid macrophages^15,16^ [Losslein, AK, Henneke, P, Ann Rev Immunol (in press)]. Granulomas are thought to limit mycobacterial dissemination, but also serve as reservoirs for mycobacterial persistence and may facilitate reactivation and spread, e.g. by necrotic transformation and promotion of tissue damage^17^.

The preference of mycobacteria for macrophages including their slow replication rate makes them a unique tool to investigate the role of *bona fide* KCs and monocytes in tissue macrophage plasticity and diversity. At the same time, the investigation of tissue-specific macrophage responses during chronic infections may provide fundamental insights into how macrophages adapt to challenges. Exploitation of the dual reporter fate-mapping mouse *Clec4f^Cre^*^-tdTomato-NLS^:*ROSA26*^EYFP^ enabled long-term tracing of KCs by multidimensional analysis, thereby uncovering the emergence of a unique, highly specialized, and adapted KC subset, which we denominated KC^low^. KC^low^ were characterized by the loss of KC identity markers, metabolic alterations, - i.e., the upregulation of inducible nitric oxide synthase (iNOS), a key enzyme in granuloma formation and antimycobacterial defense^18,19^ - and enhanced efferocytosis. These changes were accompanied by a distinct localization of KC^low^ to granuloma cores with direct mycobacterial contact, while bone marrow-derived cells induced and surrounded granulomas. Yet, KC^low^ retained unexpected plasticity and were able to regain classical immunophenotypic KC traits.

## Results

### A specific KC subset forms the core of granulomas

In order to dissect KC dynamics during mycobacterial infections with high resolution, we created a dual reporter mouse by crossing the *Clec4f^Cre^*^-tdTomato-NLS^ to a *ROSA26*^EYFP^ mouse, with tdTomato expression under control of the KC specific *Clec4f* promoter and tagged with a nuclear localisation signal (NLS)^20^ (Fig. 1a). Thus, steady-state KCs appear both tdTomato and YFP.

**Figure 1:**
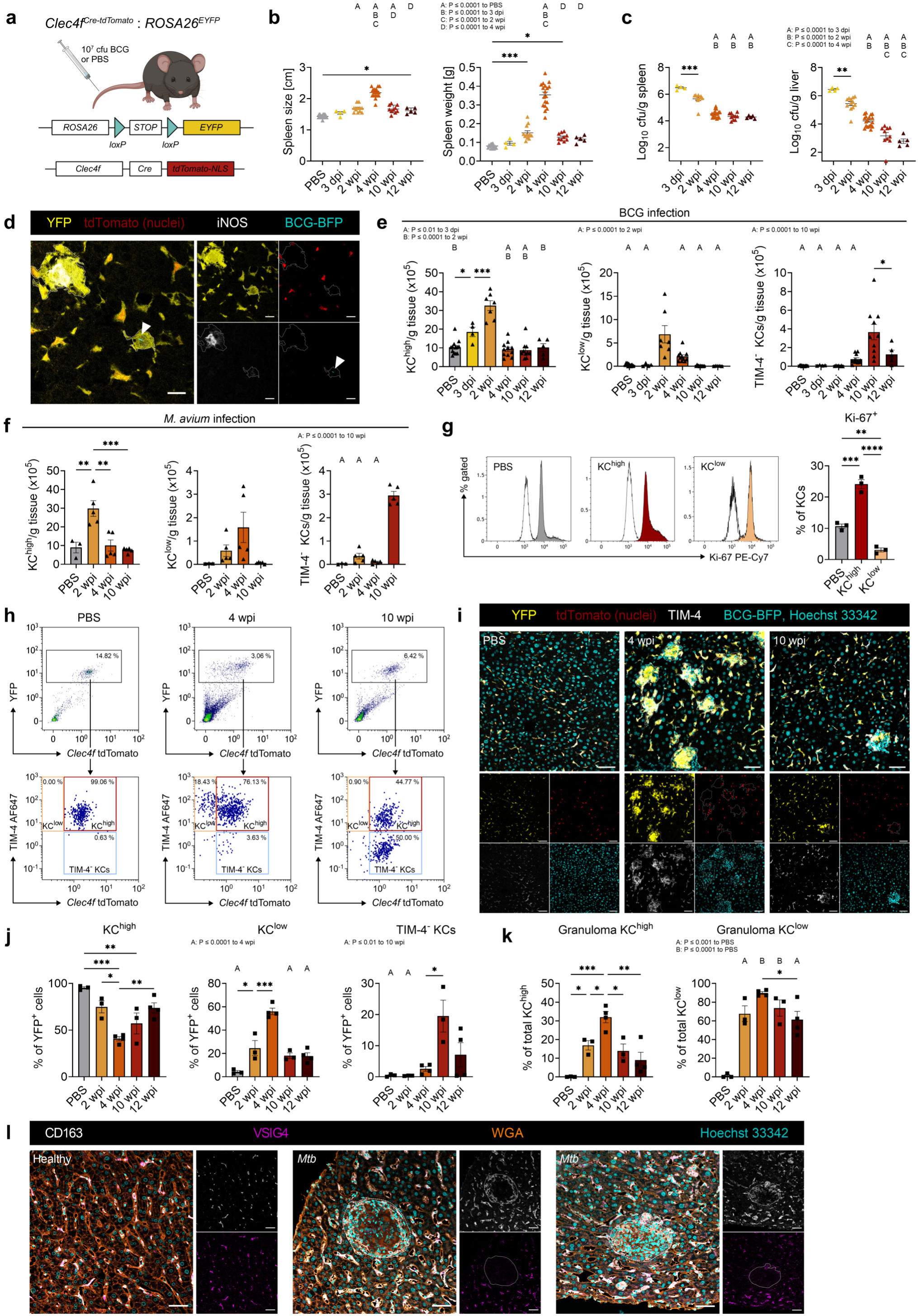
Identification of a granuloma specific Kupffer cell population (a) Experimental setup of *Clec4f^Cre-tdTomato-NLS^:ROSA26^EYFP^* mice i.v. injected with 10^7^ cfu of BCG-BFP or PBS for (b-k). (b) Spleen size and weight during infection (n=(17, 4, 12, 18, 10, 5), mice respectively). (c) Bacterial burden in spleen and liver during infection. nspleen=(4, 10, 18, 10, 5) and nliver=(4, 12, 18, 10, 5), in sequence. One mouse at 10 wpi was below the detection limit for the liver. (d) Immunofluorescence (IF) staining of a liver 4 wpi. Dotted lines mark granulomas. Arrowhead indicates BCG. Scale bars: 20 µm. (e, f) Flow cytometry of KC populations in *Clec4f^Cre-tdTomato-NLS^:ROSA26^EYFP^* mice with high (KC^high^) or low (KC^low^) tdTomato expression, or KCs lacking TIM-4 during BCG-BFP (e) or WT *M. avium* (f) infection. nBCG= (13, 4, 7, 11, 10, 5) and n*M. avium*=(3, 5, 5, 5), mice respectively. (g) Flow cytometry of Ki-67 in sorted KCs of PBS injected mice or at 2 wpi with BCG-BFP (KC^high^ and KC^low^). Isotype controls of the respective KC population are depicted in white. n=3 per group. (h) Flow cytometry gating of YFP^+^ KCs after PBS injection, 4 and 10 wpi with BCG-BFP as used in (e). Percentages of gated populations are depicted in the graph. (i) IF staining of *Clec4f^Cre-tdTomato-NLS^:ROSA26^EYFP^* livers of PBS injected mice, 4 and 10 wpi. Dotted lines mark granulomas. Scale bars: 50 µm. (j) Distribution of YFP^+^ KC subsets analysed from microscopy images as shown in (i). n=(3, 3, 4, 3, 4) in sequence. (k) Percentage of granuloma-associated KC^high^ and KC^low^ based on (i, j). (l) IF staining of human livers from a healthy donor and two patients infected with *Mycobacterium tuberculosis* (*Mtb*). Dotted lines mark granulomas. Scale bars: 50 µm. Data represent mean ± SEM with each symbol depicting one mouse are from at least 2 independent experiments. One-way ANOVA with Tukey‘s multiple comparisons test. * *P* ≤ 0.05, ** *P* ≤ 0.01, *** *P* ≤ 0.001, **** *P* ≤ 0.0001. dpi, days post infection. wpi, weeks post infection.

Intravenous infection with 10^7^ colony forming units (cfu) of *Mycobacterium (M.) bovis* Bacillus Calmette-Guerin (BCG) led to a persistent, systemic infection, associated with splenomegaly most pronounced at 4 weeks post infection (wpi) (Fig. 1b). Splenic and hepatic bacterial burdens peaked in the first week and then showed a slow, gradual decrease over several months, indicating a chronic infection (Fig. 1c). We observed a similar progression in response to *M. avium* (Extended Data Fig. 1a), the most common cause of nontuberculous infections in humans in many parts of the world^21^.

Both infection models caused formation of liver granulomas as early as 2 wpi, which decreased over time in number and size (Extended Data Fig 1b, c). Additionally, we identified mycobacteria residing in granulomas, which contained abundant iNOS-expressing macrophages, indicating an active and mycobacteria-specific inflammatory response (Fig. 1d).

Flow cytometry analysis uncovered KC diversification after infection with either BCG or *M. avium*. The PBS control group contained only tdTomato and YFP double positive *bona fide* KCs, which additionally expressed the resident tissue macrophage marker TIM-4^6^, and were termed KC^high^. Interestingly, KC^high^ increased steeply until 2 wpi, before returning to homeostatic level (Fig. 1e, f). This was in line with initial proliferation of KC^high^ (Fig. 1g), which likely served to control the infection until monocyte influx compensated for an increased macrophage need.

Notably, early in infection (2-4 wpi), a second YFP^+^ KC subset emerged, defined by low to abrogated tdTomato expression (KC^low^). In addition, TIM-4^-^ KCs were identified from 4 wpi onwards and peaked at 10 wpi (Fig. 1e, f, h).

Microscopic analysis confirmed the appearance of KC^low^, predominantly at 4 wpi (Fig. 1i, j). Yet, in contrast to the flow cytometry data, KC^low^ persisted until late infection time points (10- 12 wpi), when granuloma numbers were substantially decreased (Fig. 1i, j). In view of the relatively low number of cells, it seems probable, that the microscopy, which enabled focusing on granulomas, captured KC^low^ better than flow cytometry. On the other hand, the late microscopic emergence of TIM-4^-^ KCs accurately reflected the flow cytometry data.

*Clec4f* reporter downregulation specifically occurred for KCs in granuloma cores, while KCs surrounding granulomas remained KC^high^ (Fig. 1i). For KC^high^, the peak association to granulomas was at 4 wpi (∼32 % of total KC^high^), while up to ∼90 % of KC^low^ resided within granulomas at this infection stage (Fig. 1k). The relative association of KC^low^ with granulomas decreased until 12 wpi, but remained higher than for KC^high^ (Fig. 1k). In other words, KC^low^ rather than KC^high^ stood out as granuloma macrophages.

The maintenance of the KC phenotype and its acquisition by MoKCs respectively have been attributed to the interaction with HSCs via bone morphogenetic proteins and DLL4-Notch signaling with LSECs^4,7,8^. Therefore, we explored by microscopy whether the phenotypical changes of KC^low^ were caused by a loss of heterocellular interactions. We found a marked increase of HSCs around granulomas revealed by staining for desmin^22^ (Extended Data Fig. 2a). Moreover, KC^low^ resided partly in the immediate vicinity of HSCs and, albeit at a lower extend, LSECs, even within granulomas (Extended Data Fig. 2b, c). These observations argued against loss of heterocellular contact as the central cause for CLEC4F downregulation in infection, but rather pointed at infection-induced macrophage reprogramming.

Next, we investigated whether KC^low^ analogues in mycobacterial infections exist in humans. Notably, liver biopsies from patients with *Mycobacterium tuberculosis* (*Mtb*) infection or other mycobacterial species are rare^23^, as *Mtb* infections are usually diagnosed by lung-related clinical and microbiological findings, and the puncturing of the diseased liver poses a considerable risk. Yet, we identified samples from two patients with confirmed *Mtb* infections and liver granuloma formation (Extended Data Fig. 1d). Staining for VSIG4 and CD163 identified KCs as double positive^24,25^, both in healthy and diseased liver tissue outside granulomas. However, inside granulomas, macrophages expressed only CD163 (Fig. 1l). The lack of the prototypic KC marker VSIG4 in granuloma-associated macrophages suggested the existence of KC^low^ analogous cells in humans with *Mtb* infections.

Taken together, mycobacterial infections led to the emergence of a unique KC subset - KC^low^ - that lost key immunophenotypic markers and distinctly localized to granuloma cores with close mycobacterial contact.

### KC^low^ are derived from resident KCs

In order to explore the KC^low^ origin, we combined the intravenous BGC infection model with chimeric mice generated by transplantation of *Clec4f^Cre^*^-tdTomato-NLS^:*ROSA26*^EYFP^ bone marrow (BM) into CD45.1 recipients, or vice versa (Fig. 2a). To prevent KC damage, livers were shielded during irradiation at the expense of a mixed chimerism (Extended Data Fig. 3a). The blood chimerism of circulating monocytes was used to calculate adjusted percentages of donor-derived cells in the liver. We found donor-derived TIM-4^+^ KCs to increase late in infection (Fig. 2b). Moreover, granuloma cores were mainly formed by recipient KCs at 4 wpi, while donor cells predominantly surrounded granulomas (Fig. 2c, d and Extended Data Fig. 3b). Most of the donor cells lacked CLEC4F or TIM-4 expression, but expressed MHC-II (Extended Data Fig. 3c). In contrast, splenic granulomas contained a significant proportion of donor-derived cells, which were also the predominant cells expressing iNOS (Extended Data Fig. 3d). At 10 wpi, both resident and donor-derived cells localized to hepatic granuloma cores (Fig. 2c, d). Accordingly, mycobacterial granuloma macrophages in the liver were spatially and dynamically organized related to their origin, appearing to result from organ specific cues.

**Figure 2:**
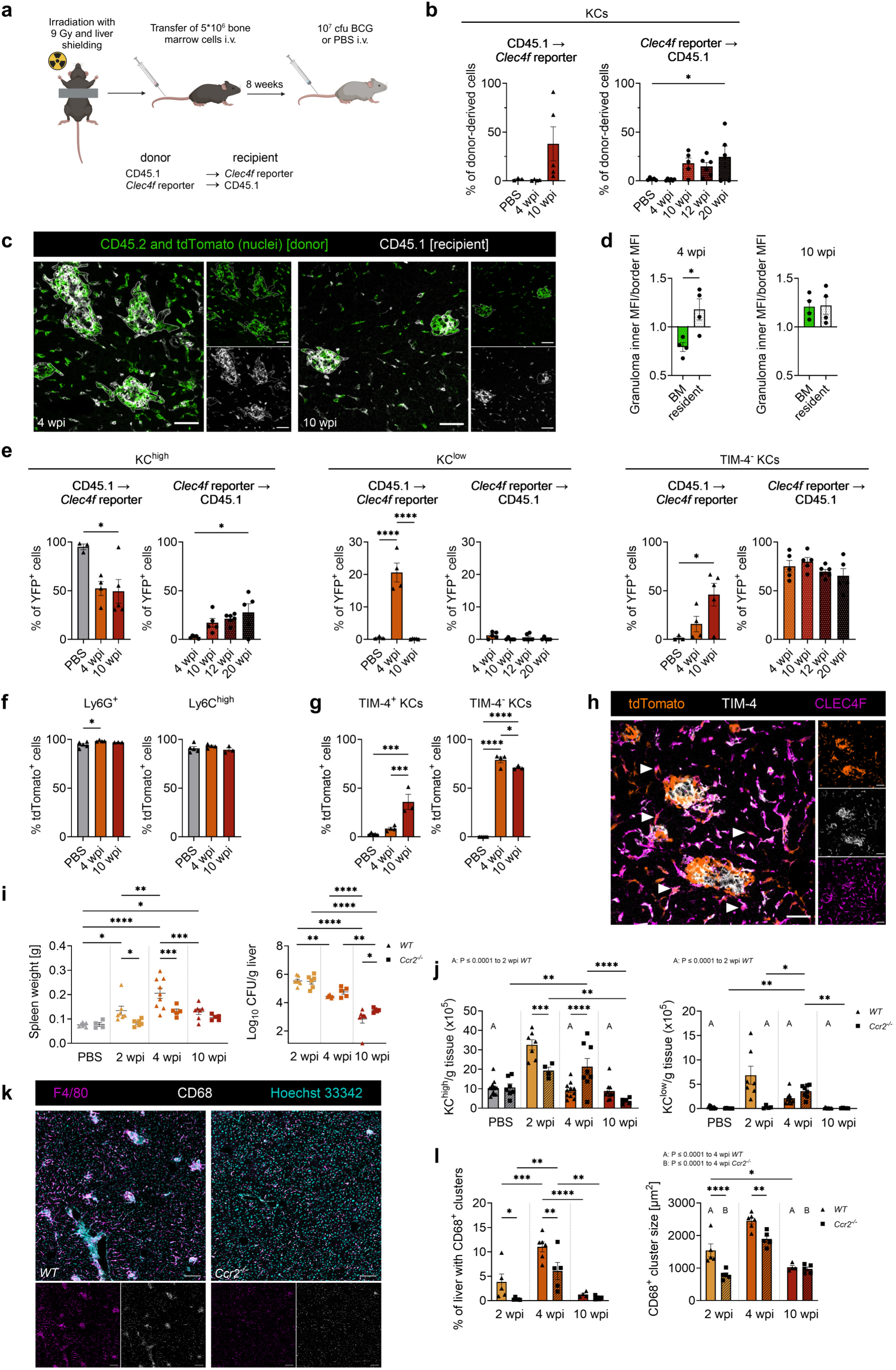
Origin of the KC subsets during infection (a) Experimental setup for (b-e). Livers were shielded from radiation and bone marrow subsequently transplanted. After 8 weeks, mice were i.v. injected with WT BCG or PBS. *Clec4f^Cre-tdTomato-NLS^:ROSA26^EYFP^* were transplanted with CD45.1 bone marrow and vice versa. (b) Chimerism of TIM-4^+^ KCs, adjusted to blood monocyte chimerism. nleft=(3, 4, 5) and nright=(5, 5, 5, 6, 5) in sequence. One-way ANOVA with Dunnett‘s multiple comparisons test compared to the PBS condition. (c) Immunofluorescence staining (IF) of a CD45.1 liver transplanted with *Clec4f^Cre-tdTomato-NLS^:ROSA26^EYFP^* bone marrow at 4 and 10 wpi. Staining against CD45.2 (donor, green) is in the same channel as the tdTomato signal. Dotted lines mark granulomas. Scale bar: 50 µm. (d) Relative mean fluorescence intensity (MFI) for CD45.2 (bone marrow (BM)-derived) or CD45.1 (resident) calculated as a ratio of liver granuloma cores and granuloma borders based on microscopic images shown in (c). Analysis was performed for 4 and 10 wpi. One dot represents averaged values for one mouse. n=4 per group. Two-tailed unpaired t test. (e) Flow cytometry analysis of KC^high^, KC^low^, and TIM-4^-^ KCs from transplanted mice in (b). (f, g) Flow cytometry analysis of tdTomato expression in Ly6G^+^ granulocytes and Ly6C^high^ monocytes (f) or in TIM-4^+^ VSIG4^+^ KCs and TIM-4^-^ VSIG4^+^ KCs (g) in livers of *Ms4a3^Cre^:ROSA26^tdTomato^* mice after BCG-BFP infection. n=(5, 5, 3), mice respectively. Two mice in the PBS condition and 4 wpi were i.v. infected via the retro-orbital route. (h) IF of a *Ms4a3^Cre^:ROSA26^tdTomato^* liver 4 wpi. Arrowheads mark exemplary tdTomato^+^ CLEC4F^+^ cells. Scale bar: 50 µm. (i) Spleen weight and hepatic bacterial burden comparing *WT* (nspleen weight=(7, 7, 9, 7), ncfu=(7, 7, 7), mice respectively) and *Ccr2^-I-^* (nspleen weight=(4, 6, 5, 5), ncfu=(7, 6, 6) in sequence) mice. Infections were performed with WT BCG. One mouse at 10 wpi was below the detection limit for hepatic bacterial burden. (j) Flow cytometry of KC^high^ and KC^low^ in *Clec4f^Cre-tdTomato-NLS^:ROSA26^EYFP^* (data from Fig. 1e) and *Clec4f^Cre-tdTomato-NLS^:ROSA26^EYFP^:Ccr2^-I-^* (n=(7, 4, 8, 5) in sequence) mice. (k) IF 2 wpi of *WT* and *Ccr2^-I-^* mouse livers. Scale bars: 100 µm. (l) Quantification of the liver area occupied by granulomas in microscopy images shown in (k), defined as CD68^+^ clusters with a minimal size of 350 µm^2^, as well as their average size. One dot represents a mouse, n*WT*=(5, 6, 4) and nCcr2-/-=(5, 5, 5) respectively. Infections were performed with WT BCG. Data represent mean ± SEM with each symbol depicting one mouse and are derived from at least 2 independent experiments. One-way ANOVA (e-g) or two-way ANOVA (i, j, l) with Tukey‘s multiple comparisons test, unless otherwise stated. * *P* ≤ 0.05, ** *P* ≤ 0.01, *** *P* ≤ 0.001, **** *P* ≤ 0.0001.

Furthermore, we noted that donor-derived KC^high^ slowly increased during prolonged infection, while donor-derived KC^low^ remained rare (Fig. 2e). In contrast, increased recipient-derived KC^low^ at 4 wpi indicated that KC^low^ derived from *bona fide* KC without substantial contribution of BM-derived cells. Additionally, and in accordance with the literature, TIM-4^-^ KCs were mainly donor-derived^6,26^. The increase of TIM-4^-^ recipient KCs likely reflected the contribution of remaining recipient BM cells due to the mixed chimerism (Fig. 2e).

To reinforce these findings, we analyzed *Ms4a3^Cre^:ROSA26^tdTomato^* fate-mapping mice, in which granulocyte-monocyte progenitor (GMP)-derived cells, and thus the vast majority of monocytes, are indefinitely labelled with tdTomato^27^. As expected, a high percentage of granulocytes and Ly6C^high^ monocytes in the liver were tdTomato-positive (Fig. 2f). In line with our expectations, tdTomato^+^ TIM-4^-^ KCs rapidly increased in number during infection, while tdTomato^+^ TIM-4^+^ KCs displayed a slow but progressive increase (Fig. 2g).

Additionally, and in full alignment with the model that mycobacterial liver granuloma cores are composed of resident KCs, microscopy of *Ms4a3^Cre^:ROSA26^tdTomato^* livers revealed a lack of tdTomato^+^, and thus monocyte-derived cells, in granuloma centers at 4 wpi (Fig. 2h). Of note, some tdTomato^+^ cell agglomerates were visible, yet, since they were negative for TIM-4, they represented immature granulomas. Moreover, we identified occasional tdTomato^+^ cells, which coexpressed CLEC4F and in part TIM-4, localized outside of granulomas, indicating a transitional stage towards KC^high^ (Fig. 2h).

Subsequently, we explored the role of monocytes during infections, given the distinct spatial distribution of BM-derived macrophages at granuloma margins 4 wpi. Accordingly, we infected *Ccr2^-I-^* mice, which are deficient in circulating Ly6C^high^ monocytes (Extended Data Fig. 3e)^28^, and found a reduced splenomegaly and increased splenic bacterial loads, while hepatic bacterial counts were comparable to *WT* mice early in infection (Fig. 2i and Extended Data Fig. 3f). Moreover, livers of *Ccr2^-I-^* mice were devoid of Ly6C^high^ monocytes during infection and showed lower granulocyte numbers as compared to *WT* mice at 2 and 4 wpi (Extended Data Fig. 3e, g).

Comparison of the KC subsets within the *Clec4f* reporter line crossed onto a *Ccr2^-I-^* background revealed reduced KC^high^ and KC^low^ numbers at 2 wpi within the knockout mice (Fig. 2j). Yet, KC^high^ were markedly increased 4 wpi in *Ccr2^-I-^ Clec4f* reporter mice, making it tempting to speculate that this increase in KC^high^ compensated for the lack of monocytes. Furthermore, at 10 wpi *Ccr2^-I-^ Clec4f* reporter mice had fewer TIM-4^-^ KCs (Extended Data Fig. 3h), in line with their BM origin. Additionally, *Ccr2^-I-^* mice displayed significantly smaller and fewer liver granulomas (Fig. 2k, l), indicating the role of monocytes in granuloma propagation and explaining the observed reduction of KC^low^ in *Ccr2^-I-^ Clec4f* reporter mice, especially at 2 wpi.

In summary, we have demonstrated that KC^low^ are a progeny of *bona fide* KCs. The role of monocytes in granuloma formation appeared to be rather indirect: they did not directly contribute to granulomas, but influenced granuloma size and counts, without substantially affecting bacterial clearance. Thus, *bona fide* KCs appeared to be the main granuloma cells, as well as key effectors for bacterial control.

### KC^low^ lose their Kupffer cell identity

To further explore the identity and function of KC^low^, we performed bulk ATAC sequencing (seq) (gating strategy in Extended Data Fig. 4a) and bulk RNAseq on the three identified KC populations at 4 wpi, and of KCs after PBS injection. The principle component analysis (PCA) of the ATACseq data, as well as analysis of the significantly differential accessible regions (DARs) revealed distinct genome-wide chromatin accessibility profiles for each subtype (Fig. 3a and Extended Data Fig. 4b). The DARs of KC^low^ were particularly close to genes involved in cellular response to low oxygen, suggesting reduced oxygen availability within granulomas. Additionally, enriched gene ontology (GO) terms of genes close to DAR of KC^low^ included actin filament and fibronectin binding (Extended Data Fig. 4c), indicating their contribution to granuloma organization. Surprisingly, enriched GO terms of genes close to DAR of TIM-4^-^ KCs were related to bacterial response (Fig. 3b), even though they appeared late in infection and were only loosely associated with granulomas.

**Figure 3:**
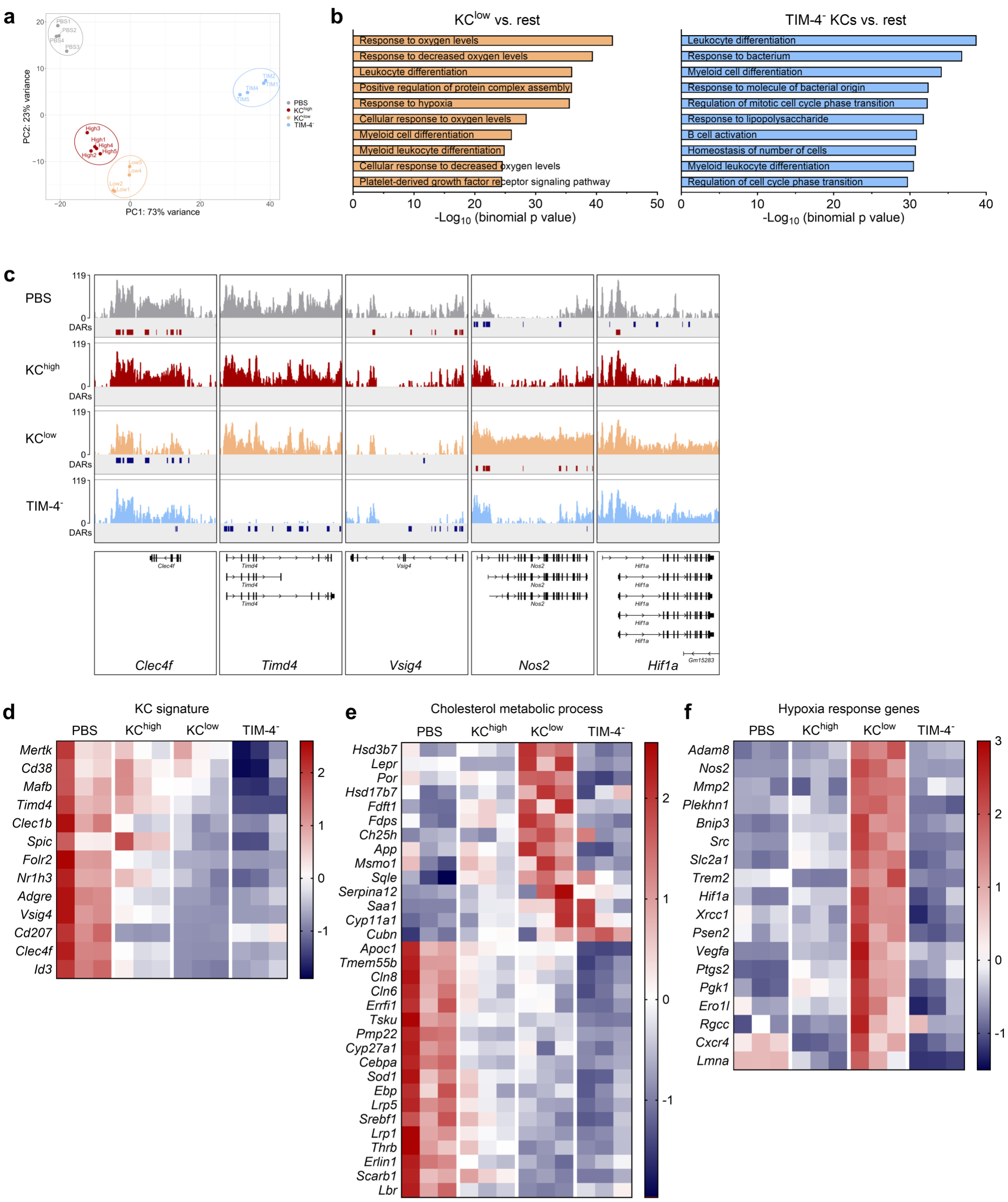
KC^low^ lose identity markers and adapt to their new microenvironmental niche (a) Principle component analysis (PCA) plot from ATAC sequencing (seq) of sorted KC^high^, KC^low^, and TIM-4^-^ KCs at 4 wpi, as well as KC^high^ from PBS treated mice. n = 5 for KC^high^ and n = 4 for the other populations from two independent infection experiments. (b) Top 10 enriched GO-terms of biological processes for KC^low^ vs. all other populations (rest) (left) and TIM-4^-^ KCs vs. rest (right) within the ATACseq data. (c) Integrative Genomics Viewer of ATACseq displaying chromatin accessibility for indicated genes representative for one sample per population. Significantly differentially accessible regions (DARs) are depicted below the tracks in blue for downregulated and red for upregulated accessibility comparing one population vs. all other populations. (d) Heatmap of Kupffer cell signature genes in bulk RNAseq data for sorted KC^high^, KC^low^, and TIM-4^-^ KCs at 4 wpi, as well as for KC^high^ from PBS treated mice. n = 3 per group. (e) Heatmap of differentially expressed genes of the GO term “cholesterol metabolic process” (GO:0008203) between PBS and KC^low^ in the bulk RNAseq data. (f) Heatmap of differentially expressed genes of the GO term “response to hypoxia” (GO:0001666) in the bulk RNAseq data.

Chromatin accessibility in the gene locus for *Clec4f* was decreased in KC^low^ and a complete absence of accessibility was observed in the *Timd4* locus for TIM-4^-^ KCs (Fig. 3c). Additionally, chromatin accessibility in KC^low^ was high for *Hif1a* and substantially increased in the *Nos2* gene (Fig. 3c). In contrast, key transcription factors of KCs^4,29^ remained largely unchanged (Extended Data Fig. 4d).

Bulk RNA sequencing uncovered distinct expression profiles of the predefined populations (Extended Data Fig. 4e). KC^low^ exhibited drastically reduced expression of many KC signature genes, most notably for *Id3*, *Clec4f*, *Cd207*, and *Vsig4* (Fig. 3d). TIM-4^-^ KCs did not yet upregulate these genes, indicating their early transitional stage towards the KC phenotype (Fig. 3d). It is worth mentioning that, although TIM-4^-^ KCs exhibited reduced *Clec4f* expression compared to KCs from PBS mice, the *Clec4f* promotor activity was sufficient to drive Cre/Lox recombination, as TIM-4^-^ KCs were identified as YFP^+^.

GO terms for KC^low^ indicated metabolic rewiring and increased chemokine receptor binding compared to KC^high^, suggesting differences in microenvironmental cues (Extended Data Fig. 4f). Mycobacteria can directly and profoundly affect the macrophage lipid metabolism^30,31^ and the accumulation of cholesterol is essential for the transformation of macrophages towards multinucleated giant cells^32^. While our mouse model did not yield MGCs, KC^low^ were in direct contact with mycobacteria, which might explain the altered gene expression pattern of the cholesterol metabolism (Fig. 3e).

In humans, the necrotic and hypoxic transformation of granulomas facilitates mycobacterial spread^33^. While *WT* mice did not show visible granuloma necrosis, in line with the literature^34,35^, KC^low^ still upregulated many genes associated to hypoxia (Fig. 3f), in accordance with GO terms resulting from ATACseq (Fig. 3b). It seems likely that oxygen availability in granuloma macrophages is altered due to vascular reorganization^36^, as indicated by increased *Vegfa* expression (Fig. 3b and Extended Data Fig. 4h).

Overall, KC^low^ displayed a fundamentally altered transcriptional profile compared to KC^high^ with more than 1500 differentially expressed genes (Extended Data Fig. 4g). In particular, KC^low^ upregulated inflammatory genes like *Nos2*, *Itgax*, and *Saa3* (Extended Data Fig. 4i), as well as genes involved in tissue remodelling [*Vegfa* and *Mmp12* (Extended Data Fig. 4h)], arguing for a pivotal role in granuloma formation and granuloma macrophage function.

In summary, KC^low^ and *bona fide* KC^high^ are distinct liver specific macrophage species with a unique immunophenotype and function as indicated by their transcriptional profiles and epigenetic traits, although they are of similar origin. Accordingly, our findings indicated previously underappreciated plasticity of KCs.

### ldentification of infection-specific KC clusters

We performed single cell RNA sequencing (scRNAseq) to precisely resolve the heterogeneity of KCs in infection on sorted KC^high^, KC^low^, TIM-4^-^ KCs, and CD11b^high^ or TIM-4^+^ cells, which contained mainly monocytes, from mice at 4 and 10 wpi, as well as corresponding populations from PBS injected control mice (sorting strategy in Extended Data Fig. 5a). Due to low cell numbers, the KC subsets were separately isolated and enriched. After removal of contaminating cells, we identified 17 distinct clusters (Fig. 4a). Based on the cell classification used for sorting, we found monocytes clustered together and separately from the KC subsets (Extended Data Fig. 5b).

**Figure 4:**
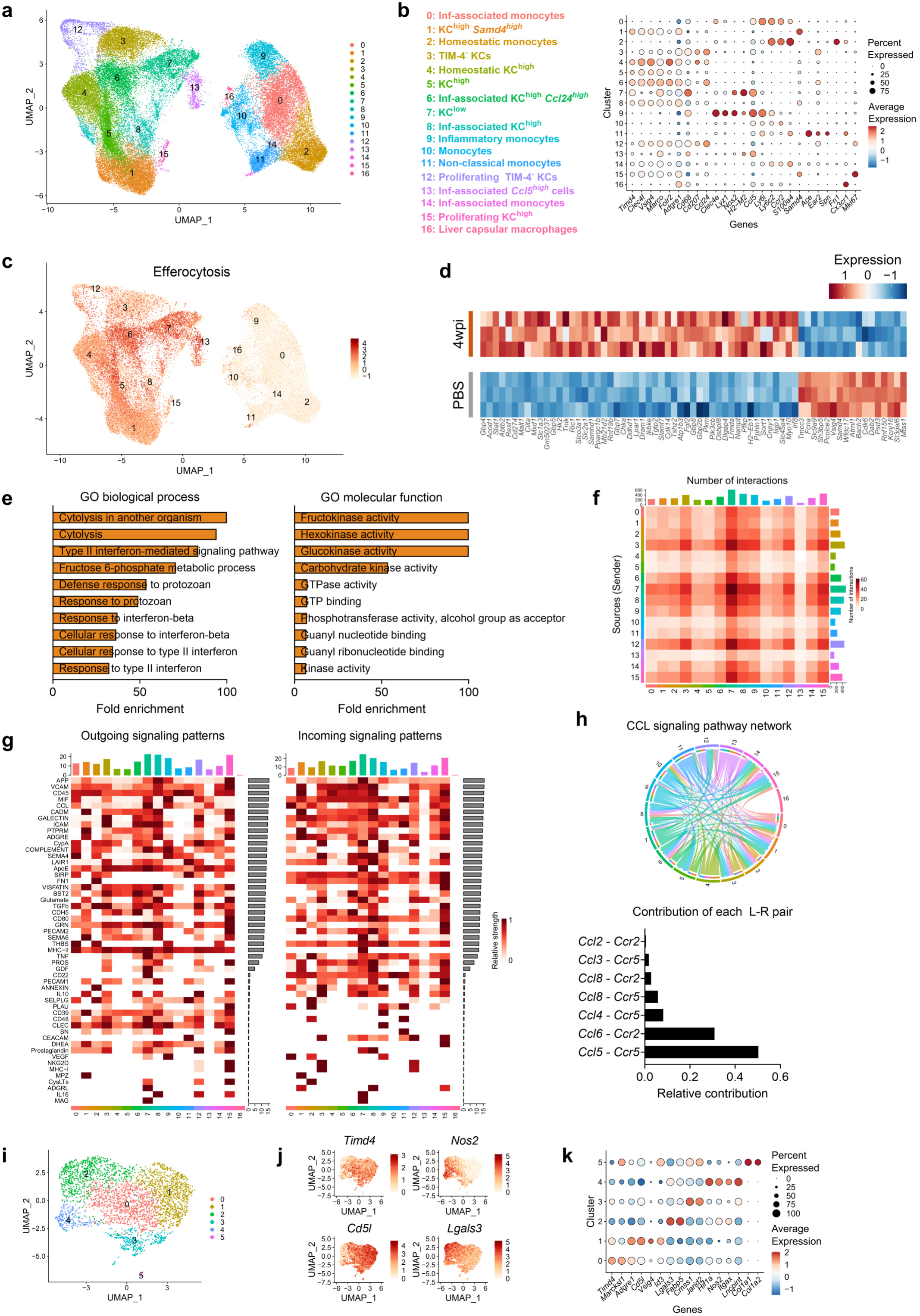
Single cell sequencing of KCs and monocytes during infection (a) Uniform manifold approximation and projection (UMAP) of sorted cells from *Clec4f^Cre-tdTomato-NLS^:ROSA26^EYFP^* mice after PBS injection or 4 and 10 weeks after BCG-BFP infection used for scRNA sequencing. In total 48,500 cells. (b) Cell cluster definition and average expression of marker genes depicted as dot plot. (c) UMAP for the aggregated gene score “efferocytosis”. (d) Heatmap of the, up to, top 50 differentially expressed genes comparing 4 wpi vs. PBS in cluster 1, calculated as pseudobulk data. (e) Top 10 GO terms in cluster 1 upregulated 4 wpi compared to the PBS control, calculated for the pseudobulk dataset. Values depicted as 100 were calculated with fold enrichment > 100. (f) Heatmap of the number of cell-cell interactions determined with CellChat exclusively for cells 4 wpi. The x-axis depicts the receivers, while the y-axis shows the clusters sending signals. Cluster 16 had no interactions and is therefore not depicted. (g) Heatmap of signalling patterns in decreased order according to the relative strength, illustrated for outgoing and incoming patterns for cells at 4 wpi. (h) Chord diagram for the CCL pathway with the interactions between the different clusters at 4 wpi, as well as the top receptor-ligand interactions contributing to the pathway. (i) Reclustered UMAP, subsetted on the KC^low^ cluster 7 from (a). In total 3183 cells. (j) Feature plots depicting the expression of the above labelled genes. (k) Dot plot for marker genes defining the different subclusters from (i).

Next, we calculated differentially expressed genes for each cluster (Extended Data Fig. 5c), thus distinguishing six KC^high^ (1, 4, 5, 6, 8, and 15), one KC^low^ (7), two TIM-4^-^ KCs (3 and 12), and six monocyte (0, 2, 9, 10, 11, and 14) clusters (Fig. 4b).

Cluster 16 represented liver capsular macrophages and cluster 13 a small KC subset, defined by high *Ccl5* expression, sorted among KC^low^ and TIM-4^-^ KCs (Fig. 4b and Extended Data Fig. 5b). The reported contribution of CCL5 as a potent chemoattractant for lymphocytes in early granuloma formation^37^ indicated an important role of cluster 13 in this context, further enhanced by cluster 7 and 9.

In accordance with our bulk RNAseq data and immunofluorescence staining, *Nos2* was highly expressed in cluster 7 (KC^low^). Additionally, monocyte cluster 9 showed increased expression of *Nos2* (Fig. 4b and Extended Data Fig. 5d). *Ccr2* expression was restricted to monocyte populations, while *Timd4* and *Clec4f*, as well as the transcriptions factors *Nr1h3* and *Spic*, characterized KC populations (Extended Data Fig. 5d). Cluster 9 potentially had a modest contribution to bacterial clearance, given the slightly elevated bacterial burden observed in *Ccr2^-I-^* mice (Fig. 2i) lacking monocytes.

Association of the clusters with the treatment modalities and time points revealed that mainly clusters 0, 3, 6, 7, 9, and 13 were enriched in infected mice, particularly at 4 wpi (Extended Data Fig. 5e, f). KC^high^ clusters 1 and 4 were primarily present in the PBS condition, indicating transcriptional changes in nearly all KCs during infection, in accordance with the bulk RNAseq data. However, these alterations seemed to be reversible, since the cluster distribution converged with the PBS condition at 10 wpi (Extended Data Fig. 5e, f).

The comparison of GO terms specific for KC^low^ (cluster 7) with those in steady state KC^high^ (cluster 0), infection-associated KC^high^ (cluster 6), or *Nos2*-expressing monocytes (cluster 9), uncovered an enrichment for endocytosis, and phagocytosis (Extended Data Fig. 5g). Additionally, KC^low^ upregulated genes associated with fatty acid synthesis and type II interferon response (Extended Data Fig. 5g), which mediates important roles in mycobacterial defense, e.g. via the induction of nitric oxide^38^.

Given that GO terms of cluster 7 were linked with phagocytosis and the recognized role of KCs in efferocytosis^39^, which has been shown to contribute to mycobacterial killing^40^ and granuloma macrophage specialization^41^, we assessed the efferocytic gene expression profile of KC^low^. The aggregated gene score for “efferocytosis” revealed an increased expression of associated genes in clusters 6, 7, and 13 (Fig. 4c).

To further investigate the differences of KC^high^ in infection, we performed pseudobulk analysis of cluster 1, comparing the PBS condition with cells isolated at 4 wpi. KC^high^ of infected mice, exhibited significant transcriptional changes (Fig. 4d), and were associated with cytolysis and type II interferon-mediated signaling (Fig. 4e). Additionally, we found an upregulated metabolic activity, including fructokinase and hexokinase (Fig. 4e). The metabolic breakdown of sugars is an important liver function^42^, and these findings point towards increased energy consumption in KCs, required to control infection.

The CellChat algorithm revealed that most interactions at 4 wpi originated from cluster 3, 7, 8, and 12, with cluster 7 notably receiving most signals (Fig. 4f, g). The top signalling patterns were associated with antigen processing and presentation (APP, MHC-II), adhesion (VCAM, ICAM), and chemokines (CCL) (Fig. 4g). Outgoing CCL signalling was present in all clusters, except for cluster 16, but most prominent in cluster 7, while infection-associated monocytes (cluster 0) received the largest input (Fig. 4g, h). This was in accordance with active monocyte recruitment to the site of infection. Among specific receptor-ligand pairs, *Ccl5* and *Ccr5* formed the top interaction, followed by *Ccl6* and *Ccr2* (Fig. 4h).

Subclustering of KC^low^ (cluster 7) to obtain a higher resolution identified six distinct subclusters (Fig. 4i and Extended Data Fig. 5h). Feature plots of *Timd4* and *Nos2* revealed discrete expression patterns, with cluster 4 exhibiting the most inflammatory profile, while expression of *Timd4* was lower than in other clusters (Fig. 4j, k). Additionally, differential expression of *Cd5l* and *Lgals3* divided the clusters (Fig. 4j, k). *Cd5l*, encoding for “apoptosis inhibitor of macrophages”, has been shown to be upregulated during *M. avium* infection, leading to increased apoptosis resistance in macrophages and may therefore facilitate mycobacterial growth^43^. On the other hand, knockout mice for *Lgals3*, encoding for galectin-3, were found to be associated with heightened susceptibility to *Mtb* infection^44,45^. This divergent expression pattern supported a model where macrophages at the infection site were functionally distinct: Cluster 2 and 4, as *Nos2* and *Hif1a* expressing clusters, were dedicated to controlling infection, while cluster 0 and 1, with a transcription profile reminiscent of KC^high^, appeared permissive for mycobacterial persistence.

### KC^low^ are able to revert to KC^high^

Since KC^low^ were characterized by a profound loss of KC identity while being polarized for essential antibacterial functions, we wondered about their fate after infection resolution when both granulomas and KC^low^ disappeared (Fig. 1e and Fig. 2k). This disappearance could be due to cell death, or a return to the classical KC^high^ phenotype, rendering previous KC^low^ undistinguishable from persisting KC^high^ or MoKCs.

RNA velocity trajectory analysis of all macrophage clusters based on the scRNAseq data identified connections at 4 wpi between cluster 7 (KC^low^) towards cluster 8 (infection-associated KC^high^). From there, additional connections existed at 10 wpi to cluster 1, suggesting that KC^low^ may revert to KC^high^-like cells in a progressive manner (Fig. 5a). Furthermore, KC^high^ cluster 5 converged towards cluster 1 in all three conditions, suggesting it as a transitional stage of KCs (Fig. 5a). In line with our previous assumption, TIM-4^-^ KCs (cluster 3, 12) showed connections towards KC^high^ (cluster 6) (Fig. 5a). Overall, the data indicated cluster 1 as a final trajectory of KCs during infection resolution and in the steady state (Extended Data Fig. 6a).

**Figure 5:**
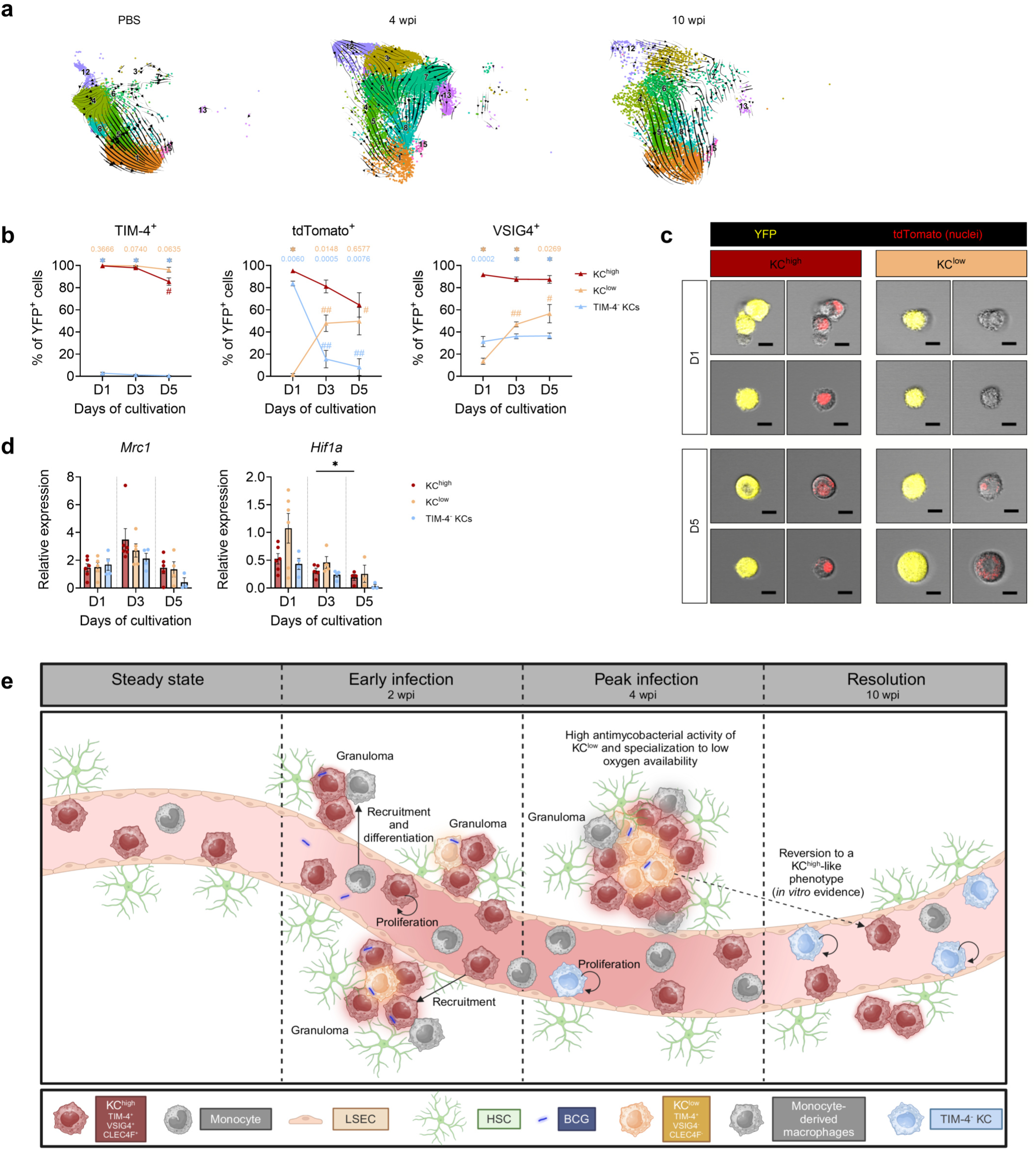
KC^low^ show remarkable plasticity (a) Trajectory analysis of the macrophage clusters contained in the scRNAseq dataset split by condition (PBS, 4 wpi, and 10 wpi), calculated based on scvelo using the dynamic model. (b) Flow cytometry of TIM-4, tdTomato, and VSIG4 expression, gated on live YFP^+^ cells, of cultured KC^high^ (n=6), KC^low^ (n=6), and TIM-4^-^ KCs (n=5 or n=4 on D5) isolated from *Clec4f^Cre-tdTomato-NLS^:ROSA26^EYFP^* livers 4 wpi with BCG-BFP. Statistical significance was calculated between a KC subset and KC^high^ at the same day (asterisks, * *P* < 0.0001 and stated p-values) and for each KC subset during the cultivation time (shown as hashtags compared to D1, # *P* ≤ 0.05, ## *P* ≤ 0.01) using a Mixed-effects model with Tukey’s multiple comparison test. (c) Confocal imaging of cultured KC^high^ and KC^low^ from (c). Depicted are two representative pictures (top and bottom) per condition and cultivation day. Bright fields are overlaid with the YFP (yellow, left) or tdTomato (red, right) reporter signal. Scale bar: 10 µm. (d) Relative gene expression of *Mrc1* and *Hif1a* normalized to *Hprt* during the cultivation of KC^high^ (n=4-6), KC^low^ (n=3-6), and Tim-4^-^ KCs (n=3-4). For *Hif1a* two values of TIM-4^-^ KCs at D5 are depicted as 0, as they were non-detectable. Mixed-effects model with Tukey’s multiple comparison test with * *P* ≤ 0.05. Each symbol depicts one mouse. (e) Schematic representation depicting the appearance and localization of the identified KC subsets during the course of infection. Data in (b-d) represent mean ± SEM and is derived from 2-3 independent experiments.

In order to corroborate these findings, we set out to analyze the KC fate *in vitro*. However, KCs are notoriously difficult to manipulate and analyze *ex vivo* since they downregulate key markers in conventional 2D cultures^46^ or in precision-cut liver sections^47^. Thus, we developed a novel system enabling *in vitro* cultivation of functional and viable KCs with preservation of KC identity parameters for several days. Our improved cultivation method involved plating sorted KCs from untreated *Clec4f^Cre^*^-tdTomato-NLS^:*ROSA26*^EYFP^ mice on Geltrex^TM^ coated wells (Extended Data Fig. 6b-d). To assess KC functionality, we analyzed their ability to avidly phagocytose BCG-BFP. Cultured KCs remained viable over 5 days with the majority of KCs continuing to express TIM- 4 (∼89 %) and maintaining *Clec4f* reporter expression (∼69 %) (Extended Data Fig. 6e, f). In contrast, both markers substantially decreased after infection with high BCG concentrations (Extended Data Fig. 6f). Moreover, VSIG4 expression was maintained with a transient decrease at D3 for the high BCG concentration (Extended Data Fig. 6f). The downregulation of both tdTomato and VSIG4 expression hinted towards the emergence of a KC^low^ analog *in vitro* in the presence of mycobacteria. KCs phagocytosed BCG over 5 days in culture (Extended Fig. 6g, h), thus demonstrating the preservation of energy consuming and therefore demanding functions of KCs in Geltrex^TM^.

Next, we sorted and cultured the predefined KC subsets extracted from infected mice. We found maintained TIM-4 expression in KC^low^, while KC^high^ exhibited a marginal decrease until day 5 (Fig. 5b). Notably, KC^low^ regained both tdTomato and VSIG4 expression over time, ultimately reaching levels comparable to KC^high^ (Fig. 5b, c). These findings suggested an unprecedented phenotypic plasticity of KCs in the absence of structural liver cells. Of note, TIM-4^-^ KCs did not increase TIM-4 expression in culture. Instead, ∼80 % of TIM-4^-^ KCs lost tdTomato expression already after one day in culture (Fig. 5b). On the transcriptional level, *Mrc1* expression, as a key hallmark across liver macrophages, was maintained over the cultivation period. Additionally, and consistent with the transformation of KC^low^ to a KC^high^-like phenotype, we observed for them a slight downregulation of *Hif1a* over time (Fig. 5d).

Overall, it seems important to note that the described *in vitro* KC culture of infected mice yielded in only a relatively small number of cells. Accordingly, although we embarked on adoptive transfers of KC^low^ we did not succeed in recovering sufficient numbers of transferred cells.

Taken together, we find that KC possess a remarkable plasticity, leading to far reaching identity loss and maximal spatial adaptations during chronic infection, yet reversion to classical *bona fide* KC features after infection clearance (Fig. 5e).

## Discussion

*Bona fide* KCs are extraordinary macrophages due to their prenatal origin, their localization in a unique microenvironment within liver sinusoids, and their frequent exposure to foreign material, e.g., to microbial components drained from the intestine. The established filtering capacity of KCs, which form the body’s largest macrophage population, manifests in their efficient phagocytic capacity, visible already minutes after intravenous bacterial injection^10,48^. The impact of the microanatomical and functional KC niche is reflected by a highly specific transcriptional profile, leading to the expression of the C-type lectin receptor CLEC4F. The dependency of KCs on the tissue environment is supported by their rapid loss in liver injuries and infections^5,49–51^, as well as the pronounced difficulty of maintaining KCs in culture^46,47^. The disappearance of *bona fide* KCs is linked with an opening of the niche and replacement by monocyte-derived macrophages, which display high plasticity during differentiation and are primed to rapidly acquire an inflammatory phenotype^5,49,52,53^.

Here we employed an infection model with *M. bovis* BCG to revisit the concept of labor division between KCs, which perform homeostatic functions, and MoKCs, which occupy newly formed niches due to tissue alterations and help to contain the spread of infection. The chronic mycobacterial infection, uncovered substantial adaptability of *bona fide* KCs, culminating in the emergence of the KC^low^ subset, characterized by the loss of lineage surface markers. Cues from their new microenvironment in granuloma cores, with a high density of mycobacteria, imposed this transformation process on KCs in both mice and humans.

Downregulation of CLEC4F in liver fibrosis or infection with *Leishmania infantum* have been described before^54,55^. Tissue reorganization and communication breakdown of KCs with HSCs and LSECs were suggested by Peiseler et al. to cause diminished expression of the lineage determining transcription factor *Nr1h3*^55^. However, we found KC^low^ to undergo their identity shift despite retaining partial close contact with HSCs and LSECs, arguing against loss of heterocellular interactions as the critical factor for the differentiation trajectory described here. It seems worth noting that *Nr1h3* expression was decreased in KC^low^ compared to KC^high^, yet, the expression levels were highly elevated compared to monocyte populations infiltrating the liver. Furthermore, ATAC sequencing revealed no significant changes in the chromatin accessibility of lineage determining KC transcription factors.

One of the inflammatory genes upregulated on KC^low^ was *ltgax* (encoding for CD11c), a marker often used to denominated dendritic cells (DCs)^56^. DCs were reported to be recruited to maturing granulomas^16^ and to phagocytose mycobacteria^57,58^. With the help of a *CD11c-eYFP* reporter mouse, Harding et al. found YFP^+^ cells, defined as DCs, as the predominant granuloma cells, able of seeding new liver granulomas^58^. In contrast, our fate-mapping investigations strongly suggest, that CD11c in granuloma cores identifies transformed *bona fide* KCs (KC^low^). The misidentification of the cells is further promoted by the upregulation of the antigen processing and presentation machinery within KC^low^. Yet, it remains to be investigated if KC^low^ are able to traffic between sites and seed granulomas, as Harding et al. proposed for the YFP^+^ cells^58^.

In addition, KC^low^ exhibited some similarities with lipid-associated macrophage-like KCs (LAM-like KCs) observed in non-infectious liver injury^59^. Both cell subsets were derived from *bona fide* KCs, show efferocytic activity, and upregulated lipid-associated genes, *Mmp12,* and *Itgax*^59^. Yet, KC^low^ are clearly distinguished from LAM-like KCs by extensive loss of both CLEC4F and VSIG4 expression, and by being maintained by differentiation of KC^high^ rather than self-proliferation.

Next to KC^low^, we also found a temporal emergence of TIM-4^-^ KCs, peaking late in infection. Data generated in *Ms4a3^Cre^:ROSA26^tdTomato^* mice indicated TIM-4^-^ KCs to be BM-derived, confirming expectations based on prior reports concerning TIM-4 upregulation on MoKCs^6^. The late appearance of this subset in infection is in accordance with the time consuming process of complete KC maturation. Corroborating this notion, we observed the highest recruitment of monocytes at 4 wpi, at which time few TIM-4^-^ KCs were present. However, only 70-80 % of TIM-4^-^ KCs in *Ms4a3^Cre^:ROSA26^tdTomato^* mice expressed tdTomato, suggesting that either a part of the population was derived from monocyte-dendritic cell progenitors (MDPs) and not GMPs, or that *bona fide* KCs downregulated TIM-4, thereby contributing to TIM-4^-^ KCs. The presence of TIM-4^-^ KCs in *Clec4f^Cre^*^-tdTomato-NLS^:*ROSA26*^EYFP^:*Ccr2^-I-^* mice argues for a downregulation of TIM-4 by a subset of KC^high^.

Interestingly, TIM-4^-^ KCs showed an epigenetic profile that resembled highly inflammatory cells, although this was not reflected on the transcriptional level, likely representing an epigenetic priming originating from BM progenitors. This prevention to develop the full inflammatory phenotype was potentially caused by insufficient contact with BCG, as TIM-4^-^ KCs had only a slight contribution to granulomas. Previous studies have identified transcriptional changes of BM progenitors in the presence of BCG, leading to an enhanced antimycobacterial capacity of monocytes, supporting our hypothesis of primed but “frustrated” TIM-4^-^ KCs^60^.

Combined with the *Ccr2^-I-^* mouse data, it seems likely that monocytes and their progeny, such as TIM-4^-^ KCs, are not essential for the initial bacterial clearance in our model. Yet, the presence of monocytes was required for granuloma induction as their lack caused a decrease in granuloma number and size consistent with reports from aerosol *Mtb* lung infections^61^. Monocytes and their precursors appear to have important roles in specific mycobacterial infection situations, in particular when the antimycobacterial capacity of resident tissue macrophages is overwhelmed, e.g., in soft tissue infections with BCG^53^, or as effector cells giving rise to specialized granuloma cells, e.g., MGCs^32^. Given our finding that mainly resident macrophages, i.e., KCs, were present in liver granuloma cores at peak infection and upregulated inflammatory markers, while the reverse was observed in the spleen, it is likely that the mycobacterial response is tissue specific. The unique reaction of liver is supported by the rarity of human hepatic mycobacterial infections, while other tissues are more commonly target of extrapulmonary *Mtb* infections^62,63^.

The return to homeostatic conditions in our model was coupled with the disappearance of KC^low^, which may indicate their inevitable cell death, caused by exhaustion after bacterial clearance and a terminal differentiation state suggested by the loss of lineage-determining markers. However, our newly established *in vitro* culture uncovered a before unknown plasticity of KCs, which retained, at least in part, the ability to regain KC signature properties. In other words, becoming KC^low^, which was associated with relocalization into granulomas and therefore the adaptation to a new microenvironment, was not a one-way road. Extraction from this inflammatory context enabled the reversion to a KC^high^-like phenotype.

Moreover, maintenance of the KC phenotype is linked to constant provision of transforming growth factor-b (TGF-β) by LSECs^8^. However, in our hands, KC marker expression was preserved in pure KC^high^ culture, strongly suggesting that sustained interactions with LSECs or HSCs are not a prerequisite for the KC identity preservation over shorter times. As TGF-β can also be produced by KCs^64–66^, autocrine signalling might have compensated for the heterocellular interaction. In contrast, TIM-4^-^ KCs failed to upregulate TIM-4 and lost their *Clec4f* reporter expression rapidly under the same conditions, indicating that they were not sufficiently primed towards the KC phenotype.

In summary, our data argues for an unprecedented plasticity of *a priori* liver-resident KCs that are both an integral part of the longitudinal diversification of the tissue macrophage landscape with integration of monocyte progeny, and the mycobacterial control, leading to the restoration of homeostasis.

## Methods

### Mice

C57BL/6J mice were purchased from Jackson Laboratories (USA) or Charles River Laboratories (Germany). *Ccr2^-I-^* (B6.129S4-*Ccr2^tm1lfc^*/J), CD45.1 (B6.SJL-*Ptprc^a^ Pepc^b^*/BoyJ or C57BL/6J-*Ptprc*^em6Lutzy^/J), *Clec4f^Cre^*^-tdTomato-NLS^ (C57BL/6J-*Clec4f^em1(cre)Glass^I*J), and *Ms4a3^Cre^* mice (C57BL/6J-Ms4a3^em2(cre)Fgnx^/J) were purchased from Jackson Laboratories (USA). For fate mapping, animals were either crossed to *ROSA26^EYFP^* (B6.129X1- *Gt(ROSA)26Sor^tm1(EYFP)Cos^I*J) or *ROSA26^tdTomato^* (B6.Cg-Gt(ROSA)26Sor^tm9(CAG-tdTomato)Hze^/J) mice, which were also purchased from Jackson Laboratories (USA).

Mice were bred in the CEMT animal facility in Freiburg, Germany, and housed under specific pathogen-free conditions with food and water *ad libitum*. Day and night cycles were set to 12h. At the start of experiments mice were typically between 6-9 weeks of age and both sexes were used. All animal experiments were approved by the Federal Ministry for Nature, Environment and Consumer’s protection of the state of Baden-WOrttemberg (proposal numbers: G19/171, G20/157, G22/076).

### Human samples

Liver biopsies were collected from individuals with diagnosed *Mycobacterium tuberculosis* infections and were provided by the BioMaterialBank Nord, Borstel, Germany. Two samples contained liver granulomas and were used for the analysis. Diagnosis was based on culture, staining and/ or PCR.

Healthy liver tissue was archived at the Institute of Surgical Pathology, Freiburg, Germany (E 324/09-121068). All experimental procedures were approved by an ethics committee at the Universitat zu LObeck (proposal number: 2024-535).

### Bacterial culture

*M. bovis* BCG was a kind gift of Prof. Dr. Dirk Wagner. BCG-BFP was generated by electroporation (2500 V, 25 µF, 1000 Ohm) of 100 µl competent BCG with 1 µg DNA of a purified BFP plasmid (Addgene plasmid # 30177; http://n2t.net/addgene:30177; RRID:Addgene_30177)^67^. Bacteria were grown in liquid culture of 7H9 broth with 10 % OADC and 0.5 % glycerol for 24 h. Afterwards, 50 µg/ml hygromycin were added for selection. Cultures were grown up to an OD600 of 1, pelleted and aliquoted in 7H9 broth with 10 % OADC and 15 % glycerol. Aliquots were stored at -80 °C till usage. *M. avium* strain 104, as well as a fluorescently labelled *M. avium* RFP were a kind gift of Prof. Dr. Trude Flo. 20 µg/ml kanamycin were added for selection of *M. avium* RFP.

### Infection

Bacterial stocks were thawed and pulled through a 27 G syringe several times to obtain a single cell solution. Additionally, stocks were thrice sonicated for 30 s followed each round by vigorous vortexing. Mice were injected i.v. into the tail vein - if not otherwise indicated - with 10^7^ cfu *M. bovis* BCG or *M. avium* in 100 µl, or received 100 µl PBS as a control injection. BCG-BFP was used for the infections unless otherwise stated.

### Bacterial burden in organs

To determine colony forming units (cfu), livers and spleens were weighed and smashed through a 70 µm filter within 500 µl PBS. Serial dilutions of the cell suspension were plated on 7H10 agar plates and incubated at 37 °C and 5 % CO2 for up to four weeks. Bacterial counts below the detection limit were depicted as half the detection limit, and used as such for statistical analysis.

### Irradiation and transplantation

For transplantations, bone marrow was isolated from CD45.1 mice or *Clec4f^Cre^*^-tdTom-^ NLS:*ROSA26EYFP* mice which were sex and age matched to the recipient. To this end, the tibia and femur from the donor mice were removed, cleaned from tissue and flushed with PBS through a 70 µm cell strainer. Cells were pelleted by centrifugation (314 g, 7 min, 4 °C).

Recipient mice were anesthetized with a ketamine-xylazine mix and livers shielded with a 2 cm wide and 1 mm thick led plate. Irradiation was performed with 9 Gray. Afterwards, 5*10^6^ isolated bone marrow cells were i.v. injected. After eight weeks, mice were injected either with PBS or BCG i.v. as described above.

To assess the liver chimerism, donor liver macrophages were normalized to the blood monocyte chimerism on the day of analysis. Data points were excluded in case the blood chimerism was below 14 %. In addition, two mice 4 wpi were excluded due to unusually low bacterial burdens.

### Tissue preparation Liver cell isolation

Mice were perfused with cold PBS, livers harvested, weighed and cut into small pieces. Livers were then digested for 45 min in a horizontal shaker at 37 °C within PBS containing 10 % FBS,

10 mg/ml Collagenase IV (Worthington), and 1 mg/ml DNase I (SigmaAldrich). The cell suspension was filtered through a 70 µm cell strainer, washed with 10 % FBS in PBS and centrifuged at 300 g, 5 min, 4 °C. The supernatant was discarded and the washing step repeated. Afterwards, red blood cell lysis was performed with 1x RBC lysis buffer (ThermoFisher) according to the manufacturer’s protocol.

Cells were incubated with CD16/32 for 10 min on ice, diluted with 1 % FBS, 2 mM EDTA in PBS (FACS buffer), and stained for 30 min at 4 °C for flow cytometry.

Cells were acquired on a Gallios (Beckmann), LSR Fortessa (BD Bioscience) flow cytometer or used for cell sorting on a MoFlo Astrios EQ (Beckman Coulter) or Aria Fusion (BD) device. Additional gating strategies can be found in Supplemental Figure 1.

Samples used for *in vitro* cultures, Ki-67 staining, for ATAC or single-cell RNA sequencing, were additionally processed by a 33 % percoll (Cytiva) after the digestion step. Samples were centrifuged for 12 min at 693 g without a break and the acceleration was set to 3. The supernatant was discarded and the pellet washed with FACS buffer. Afterwards, RBC lysis was performed and cells treated with CD16/32, before they were subjected to the antibody staining.

KC populations 2 wpi and from PBS control mice were sorted for later Ki-67 staining. Cells were subjected to fixable viability dye eFluor 450 (Invitrogen). Afterwards, they were fixed according to the manufacturer’s manual using the Foxp3/Transcription Factor Staining Buffer Set (Invitrogen), followed by staining for Ki-67 and subsequent acquisition on a LSR Fortessa (BD Bioscience) flow cytometer.

### Blood cell isolation

Blood was drawn from the eye of mice and collected in heparin tubes. Up to 100 µl blood were subjected twice to red blood cell lysis with the 1x RBC lysis buffer (ThermoFisher) according to the manufacture’s protocol. Cell suspensions were then incubated with CD16/32 for 10 min on ice and stained for flow cytometry. Gating strategies can be found in Supplemental Figure 1.

### Kupffer cell culture

KCs from untreated *Clec4f^Cre^*^-tdTomato-NLS^:*ROSA26*^EYFP^ mice, or KC^low^, KC^high^, and TIM-4^-^ KCs from mice 4 wpi were isolated and sorted as described above and used for culture experiments. 96-well U-bottom plates (Falcon) were coated with 75 µl of 1:200 diluted Geltrex^TM^ Reduced Growth Factor Basement Membrane Matrix (ThermoFisher) within Dulbecco’s Modified Eagle Medium (DMEM) containing GlutaMAX (ThermoFisher). The matrix was hardened for 2 h at 37 °C and 5 % CO2 and afterwards washed with 100 µl DMEM. Per well 20.000 cells within 200 µl medium (DMEM with 10 % FBS and 0.5 % ciprofloxacin) were plated and supplemented with 20 ng/ml M-CSF (PeproTech). Steady-state KCs were additionally treated with BCG-BFP at an MOI of 1 or 10, or left untreated. The cells were incubated at 37 °C and 5 % CO2 for up to 5 days and harvested for analysis by incubating with accutase (SigmaAldrich) for 10 min at 37 °C, followed by two times washing with FACS buffer. The cell suspension was incubated with CD16/32 for 10 min on ice, and stained with VSIG4 PE-Cy7, TIM-4 AF647, and CD45 PerCP-Cy5.5 for 30 min at 4 °C. Afterwards, steady-state KCs were stained with fixable viability dye eFluor 780 (ThermoFisher) for 20 min before acquisition on a LSR Fortessa (BD Bioscience). KCs isolated from mice after infection were stained shortly before the acquisition with DAPI (BioLegend) instead.

Confocal images of KC cultures grown on 8 well Ibidi slides with 40.000 cells per well were taken on a LSM880 with a 20x (N.A. 0.8) objective.

Additionally, for transcriptomic assessment, the cells were resuspended in RNA lysis buffer (QIAGEN RNeasy Micro kit^TM^) containing 1 % β-mercaptoethanol after harvesting with accutase. RNA was isolated according to the manufacturer’s instruction of the QIAGEN RNeasy Plus Micro kit^TM^.

### qPCR

Isolated RNA was reverse transcribed to cDNA using the SuperScript^TM^ IV VILO^TM^ Master Mix (Thermo Fisher) according to the manufacturer’s instruction. For quantitative PCR 5 µl of Absolute qPCR SYBR Green Mix (Thermo Fisher) were added to 0.05 µl of each primer, 2.9 µl water, and 2 µl cDNA. Samples were run on 384 well plates using a Roche Lightcycler^TM^. Relative gene expression was normalized to the expression of hypoxanthine-guanine phosphoribosyltransferase 1 (*Hprt1)*. Primer sequences are listed in table 1 below.

**Table 1:**
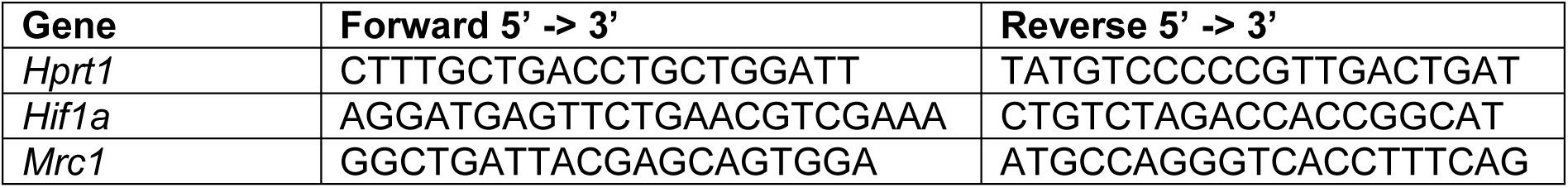
Primer sequences.

### Histological analysis

Livers and spleens were removed and fixed in 4 % PFA at 4 °C for 4 h. The samples were then transferred to 20 % sucrose till they sunk to the bottom of the tube and flash frozen with TissueTek (Sakura) in liquid nitrogen.

Fixed organs were cut on a cryotome at 8 or 30 µm thickness at -20 °C. Sections were permeabilized with wash buffer (PBS containing 1 % BSA, 0.1 % Triton-X100) at 4 °C for 5-8 h. Afterwards, blocking with CD16/32 was performed for 30 min at 4 °C. The tissue slices were then incubated with antibodies and Hoechst 33342 at 4 °C overnight. Samples were washed three times with wash buffer, after which secondary antibodies were added for 1 h at room temperature. After additional washing, samples were mounted with ProLong^TM^ Diamond Antifade Mountant (Invitrogen) or with SlowFade^TM^ Diamond Antifade Mountant (Invitrogen) if used for 3D reconstruction.

Sections were acquired on a Zeiss LSM710, LSM880, or LSM980 confocal microscope with a 20x objective (N.A. 0.8). Analysis was performed with ZEN (Zeiss, version 2012) and Fiji (version 1.53t). For 3D reconstructions, performed in IMARIS (Oxford Instruments Group, version 10.1.1), confocal images were taken on a Zeiss LSM880 with a 63x (N.A. 1.4, oil) objective.

### Microscopic analysis of KC subsets

Livers of *Clec4f^Cre^*^-tdTomato-NLS^:*ROSA26^EYFP^* mice were stained with TIM-4 AF647 and Hoechst 33342. Granulomas were outlined, and YFP^+^ cells with Hoechst 33342 nuclei counted, both outside and within granulomas. Based on the expression of TIM-4 and tdTomato cells were assigned to the KC subsets. At least 9 pictures per mouse were evaluated.

### Assessment of granuloma counts in the liver

For granuloma assessments, slides were stained with CD68 AF647, F4/80 AF488, and Hoechst 33342. In addition, sections 10 wpi were quenched with TrueBlack (Biotium) according to the manufacturer’s protocol before mounting. For each mouse three 8 µm thick sections, with at least 120 µm distance between them, were acquired as a tile scan with 5 % overlap on a LSM880 with a 20x (N.A. 0.8) objective. The pictures were stitched using the Hoechst 33342 signal in ZEN and analyzed with Fiji. The area of the liver was determined and a threshold for the CD68 signal was set. Afterwards, cell clusters were automatically determined with the “analyze particle” function and the settings for a minimal size of 350 µm^2^, as well as the inclusion of holes. The three sections were averaged for each mouse to represent a single value.

### Microscopic analysis of cell origin in granulomas

For the assessment of cell origin in granulomas after BCG infection, sections from CD45.1 irradiated mice transplanted with CD45.2 (*Clec4f^Cre^*^-tdTomato-NLS^:*ROSA26^EYFP^*) bone marrow and liver shielding were stained with CD45.1 APC, CD45.2 PE and Hoechst 33342. Single snap shots on an LSM880 with a 20x objective were taken in a blinded fashion by only using the Hoechst 33342 channel during acquisition. Analysis was performed in ZEN with the profile function. To be unbiased, the channels for CD45.1 and CD45.2 were set to the same color and two intersecting lines per granuloma were drawn to determine the MFI for the length of these lines. The first and last 15 µm of the lines were defined as “border”, while the rest was considered “inner” granuloma. The two measurements were averaged, and to obtain the ratio between inner and border granuloma, the values were divided by each other and represented as the relative MFI. MFI values of 15 granulomas were averaged for one mouse 4 wpi, while at least 10 granulomas per mouse were assessed for 10 wpi.

### Microscopic analysis of iNOS expression in the spleen

Spleen sections from CD45.1 irradiated mice transplanted with CD45.2 (*Clec4f^Cre^*^-tdTomato-NLS^:*ROSA26^EYFP^*) bone marrow and liver shielding were stained with CD45.1 e450, CD45.2 PE, CD68 AF647 and iNOS AF488. 10 pictures per mouse were taken on an LSM880 with a 20x objective and analyzed in Fiji. Areas of iNOS, CD45.2, and CD45.1 were determined by setting a fixed threshold for each channel. Then the overlap between iNOS and CD45.2 or iNOS and CD45.1 were evaluated with the “AND” function. Data was depicted as percentage of CD45.1 or CD45.2 overlapping iNOS area compared to the total iNOS area.

### Multicycle immunofluorescence staining

For multicycle IF stainings, slides were blocked for 1 h at room temperature with wash buffer and then stained for 3 h at room temperature. The antibodies per cycle can be found in table 2. Every staining round Hoechst 33342 was included to have a reference for merging the pictures later on. After staining, samples were washed three times with wash buffer and mounted with SlowFade^TM^ Diamond Antifade Mountant (Thermo Fisher).

**Table 2:**
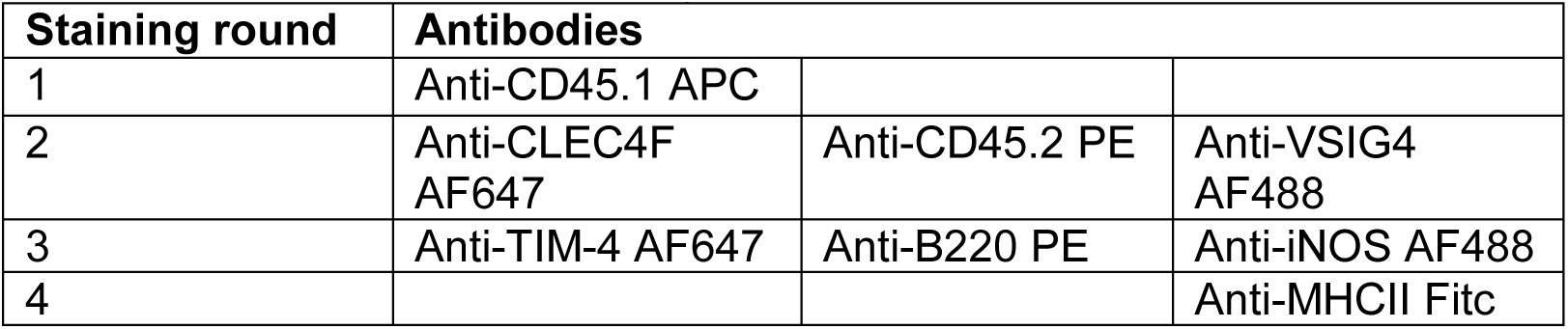
Antibodies used for multicycle IF.

| Staining round | Antibodies |  |  |
| --- | --- | --- | --- |
| 1 | Anti-CD45.1 APC |  |  |
| 2 | Anti-CLEC4F AF647 | Anti-CD45.2 PE | Anti-VSIG4 AF488 |
| 3 | Anti-TIM-4 AF647 | Anti-B220 PE | Anti-iNOS AF488 |
| 4 |  |  | Anti-MHCII Fitc |

Sections were acquired on a Zeiss LSM980 confocal microscope with a 20x objective (N.A. 0.8).

Afterwards, slides were demounted in PBS and destained for 20 min at room temperature with 0.002 % SDS, 20 mM NaOH in H2O under rocking and light exposure. Sections were then four times washed with H2O, mounted and acquired on the microscope as a reference for the background noise. Slides, were then subjected to a new round of staining and destaining. Pictures were analyzed with ZEN.

### Histological preparation of human samples

Paraffin embedded patient liver samples were cut with 2 µm thickness, dried and de- paraffinized. Antigen retrieval was performed in citrate buffer for 5 min. Afterwards, the tissue was blocked in 5 % BSA in PBS for 30 min and then incubated with the primary antibodies against CD163 and VSIG4 over night at 4 °C. Antibodies were diluted in Antibody Diluent with background reducing components (Agilent Dako). Sections were washed with 1:10 diluted Dako Wash Buffer 10x (Agilent Dako) and then secondary antibodies, wheat germ agglutinin (WGA), and Hoechst 33342 were added for 45 min. Sections were washed again and mounted with ProLong^TM^ Gold Antifade (Invitrogen) mounting medium. Additionally, serial sections were used for a hematoxylin and eosin (H&E) staining. Images were acquired on a Zeiss LSM880 confocal microscope or an Axioscan 7 (Zeiss) with a 20x objective. Analysis was performed with ZEN.

### Bulk RNA sequencing

For bulk RNA sequencing KCs from three PBS injected mice and KC^low^, KC^high^, and TIM-4^-^ KCs of three mice 4 wpi were directly sorted into RNA lysis buffer containing 1 % β- mercaptoethanol. RNA was isolated with the ExtractMe Total RNA Micro Spin Kit (Blirt) according to the manufacturer’s protocol.

Sequencing was performed by CeGaT (TObingen, Germany) using the NovaSeq 6000 (Illumina) system with 2x 100 bp read length with 50 million clusters/sample. The company provided trimmed reads (done with STAR version 2.7.3) which were aligned to the reference genome mm10. The data was analyzed using DESeq2 (version 1.40.2) in R (version 4.3.2)^68^. Genes with row sums below 2 were excluded. The PCA plot was produced by the plotPCA function in R. All heatmaps depict calculated Z values from normalized reads. Volcano plots were prepared in Prism^TM^ 10 (GraphPad) by depicting log2 fold changes and the adjusted p values (padj), for which the Benjamini-Hochberg adjustment was used. Genes with a padj < 0.05 were considered as differentially expressed and marked in a color for log2 fold change > 2 or < -2. GO terms for differentially expressed genes with padj < 0.05, were determined with PANTHER (version 19.0, https://www.pantherdb.org/).

### Assay for transposase-accessible chromatin (ATAC) sequencing

ATACseq was performed on liver macrophages from five mice 4 wpi, and four PBS control mice. The KC^low^ and TIM-4^-^ population from one infected mouse were excluded due to a technical issue. Sorted KCs were frozen in RPMI containing 10 % DMSO with 10 % FBS. After thawing at 37°C, samples were centrifuged (500 g, 5 min, 4 °C) and resuspended in 250 µl OMNI buffer (10 mM Tris-HCl (pH 7.5), 10 mM NaCl, 3 mM MgCl2, 0.1 % IGEPAL-CA630 (SigmaAldrich), 0.1 % Tween-20, 0.01 % Digitonin (Promega) in DMSO). After 5 min of permeabilization, nuclei were centrifuged (500 g, 5 min, 4 °C) and the supernatant discarded. 180 µl tagmentation buffer (33 mM Tris-acetate, 66 mM K-acetate (SigmaAldrich), 11 mM Mg-acetate, 16 % *N,N*-Dimethylformamide (EMD Millipore)) was added without disturbing the pellet. After centrifugation (500 g, 5 min, 4 °C) the nuclei were resuspended in 15 µl cold tagmentation buffer, counted, and the concentration adjusted. 1 µl Tn5 Illumina Tagment DNA Enzyme (Illumina) was added to 20 µl nuclei suspension and incubated for 60 min with 500 rpm at 37 °C. Tagmented DNA was purified using the MinElute Reaction Cleanup Kit (Qiagen). DNA fragment amplification was performed by addition of 25 µl NEBNext High-Fidelity 2x PCR MasterMix (New England Biolabs) to 10 µl tagmented DNA, 0.7 µl 24 UDI for Tagmented libraries – Set I (Diagenode) and 14.3 µl Molecular Biology Grade Water (Corning) using 8 PCR-cycles. Amplified DNA was purified with the MinElute Reaction Cleanup Kit (Qiagen). Very small and large fragments were removed using a double-sided bead clean-up with a right side clean-up ratio of 0.55X and a left side clean-up ratio of 1.5X (AMPure XP beads, Beckmann Coulter). Fragment size distribution of final libraries was assessed with a TapeStation (Agilent). Libraries were sequenced on a NextSeq 1000 (Illumina).

Processing of the sequencing data was performed with Galaxy Europe (https://usegalaxy.eu). First, adapters were trimmed with Cutadapt (version 4.0) and the sequences aligned to the reference genome mm10 with Bowtie2 (version 2.4.5). Properly paired reads with MAPQ ≥ 30 were kept for analysis. Mitochondrial reads and reads falling into blacklisted genomic regions for mm10 defined by ENCODE^69^ were excluded. Duplicates were removed with Picard’s MarkDuplicates (version 2.18.2.3) tool with a lenient validation stringency. Peaks were called using MACS2 (version 2.2.7.1). Downstream analysis was performed in R (version 4.3.2) with DESeq2 (version 1.40.2) and batch correction, adjusting the function with “design = ∼ batch + group”. Genes with row sums below 10 were excluded. Chromatin accessible regions were depicted with the Integrative Genomics Viewer^70^ (version 2.16.2).

To assess the associated genomic regions and generate gene ontology terms, the online tool GREAT (version 4.0.4, https://great.stanford.edu/public/html/) was used with the species assembly mm10 for genes with padj < 0.05 and a log2 FoldChange > 0.5. The settings were “basal plus extension” with 5 kbp upstream, 1 kbp downstream, and a maximal extension of 1000 kbp. To contrast a single group vs. all others and derive gene ontology terms, DESeq2 was rerun with the new definition of groups. Volcano plots were prepared in Prism^TM^ 10 (GraphPad) by depicting log2 fold changes and the adjusted p values (padj), for which the Benjamini-Hochberg adjustment was used. Genes with a padj < 0.05 were considered as differentially expressed and marked in a color for log2 fold change > 2 or < -2.

Peak visualization was created using the Integrative Genomics Viewer. Significantly differentially accessible regions were calculated with DESeq2 for a single group vs. all others and depicted in blue for significantly downregulated and in red for significantly upregulated accessible regions.

### Single-cell RNA sequencing

For the single-cell RNA sequencing (scRNAseq), liver cells of three mice per group (PBS, 4 or 10 weeks after infection) were prepared as described above. During the antibody staining step, each sample was additionally incubated by a specific TotalSeq^TM^ hashtag antibody (BioLegend), contained in table 3, to allow for multiplexing.

**Table 3:** Hashtag antibodies per sample.

| Hashtag antibody | Mouse condition |
| --- | --- |
| C0304 | Mouse1 PBS |
| C0309 | Mouse2 PBS |
| C0310 | Mouse3 PBS |
| C0305 | Mouse1 4 wpi |
| C0307 | Mouse2 4 wpi |
| C0308 | Mouse3 4 wpi |
| C0301 | Mouse1 10 wpi |
| C0302 | Mouse2 10 wpi |
| C0303 | Mouse3 10 wpi |

Cells were sorted according to the strategy in Extended Figure 5A and afterwards 30.000-100.000 cells of the same population from different mice were pooled. For the TIM-4^-^ KC population 4 wpi one mouse was excluded due to low cell numbers.

ScRNAseq was performed using the Chromium Next GEM Single Cell 5’ Kit v2, and libraries were constructed with the Library Construction Kit by 10X Genomics. Samples were sequenced on a NovaSeq6000 (Illumina). Fastq files were aligned with the CellRanger software (7.2.0) with prebuilt mouse mm10 reference and downstream analysis was carried out in R (version 4.3.2) with the Seurat package (version 5.0.1)^71^. For demultiplexing, HTODemux with a positive quantile threshold of 0.99 was employed and only singlets were kept for further processing. Cells with more than 5 % mitochondrial transcripts, and less than 200 or more than 5000 detectable genes were excluded. The datasets from the different samples were subsequently merged, and subjected to the NormalizeData(),

FindVariableFeatures(), ScaleData(), and RunPCA() algorithm with default parameters. For correction of batch effects, the data was integrated with harmony^72^. FindNeighbours and FindClusters were applied with the top 30 harmony dimensions (determined by the ElbowPlot() function) and a resolution of 0.6. To depict the data RunUMAP() was calculated for the harmony reduction and plotted with DimPlot. Based on the clustering and gene expression patterns, contaminating clusters that did not contain macrophages or monocytes were excluded. The total remaining 48,500 cells were reclustered with the functions FindNeighbours and FindClusters, using the top 30 harmony dimensions and a resolution of 0.7. In addition, cluster 7 was further subsetted and reclustered using the top 30 harmony dimensions and a resolution of 0.3.

For the aggregated counts of the term “efferocytosis”, the function AddModuleScore() with the genes in Table 4 was used, identified with Mouse Genome Informatics (https://www.informatics.jax.org/) and present in the scRNAseq data.

**Table 4:**
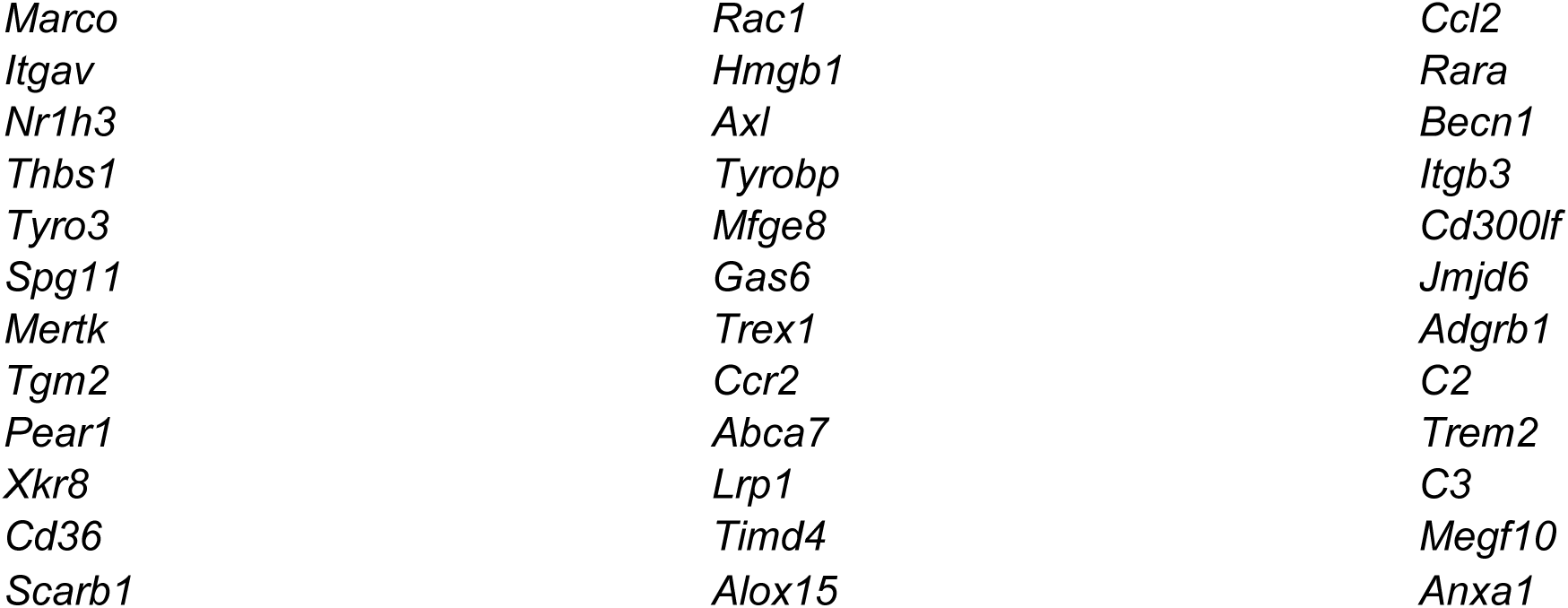
Genes used for the term efferocytosis.

Differentially expressed genes were determined with the FindAllMarkers() function. Pathway enrichment analysis was performed with DEenrichRPlot including a maximum of 2000 genes comparing cluster 7 to either cluster 0, 6 or 9. Otherwise the standard settings were used and the enrich.database was set to GO_Biological_Process_2023.

Cell communication was determined with the CellChat package (version 2.1.1), separated for the controls, 4 wpi, and 10 wpi samples. The functions identifyOverExpressedGenes(), identifyOverExpressedInteractions(), and computeCommunProb() were employed, flowed by a filtering using filterCommunication() with a minimum of 10 cells. computeCommunProbPathway() was calculated before using aggregateNet() and depiction of the data in a heatmap or chord diagram.

For pseudobulk analysis, AggregateExpression was calculated for PBS and 4 wpi samples of cluster 1. FindMarkers with DESeq2 was used to determine differentially expressed genes. GO terms for biological process and molecular function were determined with PANTHER (version 19.0, https://www.pantherdb.org/) for genes with padj < 0.05. Heatmap was plotted with the 50 most upregulated and downregulated genes with padj < 0.05.

RNA velocity was calculated using the scVelo (0.3.2) and scanpy (1.9.8) packages in Python (3.12.2)^73^. In a first step, the data was filtered and normalized, genes that did not have at least 20 counts for the spliced and unspliced layer were excluded. The top 2000 variable features were picked and further processed with the function sc.pp.neighbors() and scv.pp.moments(). For the trajectory analysis the dynamical model was calculated with scv.tl.recover_dynamics()and scv.tl.velocity for the macrophage clusters (clusters 1, 3, 4, 5, 6, 7, 8, 12, 13, 15) individually in each condition. To avoid confounding effects by the different batches, mouse 1 in the PBS condition was excluded.

### Schematics and figures

Schematics were created in BioRender.com (Henneke, P. (2025): https://BioRender.com/y64d004 and https://BioRender.com/c51o935) and figures were created using both Prism^TM^ 10 (GraphPad) and Inkscape (version 1.3.2).

### Statistical analysis

Statistical analysis was performed in Prism^TM^ 10 (GraphPad). The usage of the statistical test is indicated in the figure legends. Differences were considered as significant if p-values were ≤ 0.05. ns, not significant, * *P* ≤ 0.05, ** *P* ≤ 0.01, *** *P* ≤ 0.001, **** *P* ≤ 0.0001, unless otherwise indicated. Data is depicted as mean ± SEM unless otherwise specified.

### Antibodies

**Table 5:** Anti-mouse antibodies used in this study.

| Reagent or Resource | Source | Identifier |
| --- | --- | --- |
| Anti-CD3 APC (clone 145-2C11) | BioLegend | Cat# 100312; RRID: AB_312677 |
| Anti-CD4 PerCP-Cy5.5 (clone RM4-5) | Invitrogen | Cat# 45-0042-82; RRID: AB_1107001 |
| Anti-CD11b PE-Cy7 (clone M1/70) | Invitrogen | Cat# 25-0112-82; RRID: AB_469588 |
| Anti-CD16/CD32 (clone 93) | BioLegend | Cat# 101302; RRID: AB_312801 |
| Anti-CD19 biotin (clone 6D5) | Miltenyi Biotec | Cat# 130-119-797; RRID: AB_2801733 |
| Anti-CD31 AF647 (clone MEC13.3) | BioLegend | Cat# 102515; RRID: AB_2161029 |
| Anti-CD45 eF450 (clone 30-F11) | Invitrogen | Cat# 48-0451-82; RRID: AB_1518806 |
| Anti-CD45 FITC (clone 30-F11) | Invitrogen | Cat# 11-0451-82; RRID: AB_465050 |
| Anti-CD45 PerCP-Cy5.5 (clone 30-F11) | Invitrogen | Cat# 45-0451-82; RRID: AB_1107002 |
| Anti-CD45.1 APC (clone A20) | Invitrogen | Cat# 17-0453-81; RRID: AB_469398 |
| Anti-CD45.1 eF450 (clone A20) | Invitrogen | Cat# 48-0453-80; RRID: AB_1272189 |
| Anti-CD45.2 PE (clone 104) | BioLegend | Cat# 109807; RRID: AB_313444 |
| Anti-CD45.2 PerCP-Cy5.5 (clone 104) | Invitrogen | Cat# 45-0454-82; RRID: AB_953590 |
| Anti-CD45R (B220) PE (clone RA3-6B2) | Invitrogen | Cat# 12-0452-82; RRID: AB_465671 |
| Anti-CD64 PerCP-Cy5.5 (clone X54-5/7.1) | BioLegend | Cat# 139308; RRID: AB_2561963 |
| Anti-CD68 AF647 (clone FA-11) | BioLegend | Cat# 137004; RRID: AB_2044002 |
| Anti-CD115 APC (clone AFS98) | Miltenyi Biotec | Cat# 130-102-962; RRID: AB_2660276 |
| Anti-Clec4F AF647 (clone 3E3F9) | BioLegend | Cat# 156804; RRID: AB_2814082 |
| Anti-F4/80 AF488 (clone Cl:A3-1) | BioRad | Cat# MCA497A488T; RRID: AB_1102554 |
| Anti-iNOS AF488 (clone CXNFT) | Invitrogen | Cat# 53-5920-82; RRID: AB_2574423 |
| Anti-iNOS APC (clone CXNFT) | Invitrogen | Cat# 17-5920-80; RRID: AB_2573244 |
| Anti-Ki-67 PE-Cy7 (clone SolA15) | Invitrogen | Cat# 25-5698-82; RRID: AB_11220070 |
| Anti-Ly6C BV605 (clone HK1.4) | BioLegend | Cat# 128036; RRID: AB_2562353 |
| Anti-Ly6C PerCP-Cy5.5 (clone HK1.4) | Invitrogen | Cat# 45-5932-80; RRID: AB_2723343 |
| Anti-Ly6G biotin (clone 1A8) | BioLegend | Cat# 127604; RRID: AB_1186108 |
| Anti-Ly6G BV605 (clone 1A8) | BioLegend | Cat# 127639; RRID: AB_2565880 |
| Anti-Ly6G FITC (clone 1A8) | BD | Cat# 561105; RRID: AB_394207 |
| Anti-MHCII FITC (M5/114.15.2) | Invitrogen | Cat# 11-5321-82; RRID: AB_465232 |
| Anti-SiglecF biotin (clone ES22-10D8) | Miltenyi Biotec | Cat# 130-119-136; RRID: AB_2733492 |
| Anti-TIM-4 AF647 (clone RMT4-54) | BioLegend | Cat# 130008; RRID: AB_2271648 |
| Anti-VSIG4 PE-Cy7 (clone NLA14) | Invitrogen | Cat# 25-5752-82; RRID: AB_2637431 |
| Anti-Streptavidin eF450 | Invitrogen | Cat# 48-4317-82; RRID: AB_10359737 |
| Anti-Streptavidin PE-Cy7 | Invitrogen | Cat# 25-4317-82; RRID: AB_10116480 |
| Goat anti-CLEC4F | Invitrogen | Cat# PA5-47396; RRID: AB_2608299 |
| Goat anti-rabbit AF633 | Invitrogen | Cat# A-21071; RRID: AB_2535732 |
| Rat IgG2a kappa Isotype Control (clone eBR2a) PE-Cy7 | Invitrogen | Cat# 25-4321-82; RRID: AB_470200 |
| Rabbit anti-desmin | Invitrogen | Cat# PA5-16705; RRID: AB_10977258 |
| Rabbit anti-goat AF488 | Abcam | Cat# ab150141 |
| TotalSeq-C0304 anti-mouse Hashtag 4 (M1/42; 30-F11) | BioLegend | Cat# 155867; RRID: AB_2800696 |
| TotalSeq-C0309 anti-mouse Hashtag 9 (M1/42; 30-F11) | BioLegend | Cat# 155847; RRID: AB_2814075 |
| TotalSeq-C0310 anti-mouse Hashtag 10 (M1/42; 30-F11) | BioLegend | Cat# 155879; RRID: AB_2819914 |
| TotalSeq-C0305 anti-mouse Hashtag 5 (M1/42; 30-F11) | BioLegend | Cat# 155869; RRID: AB_2800697 |
| TotalSeq-C0307 anti-mouse Hashtag 7 (M1/42; 30-F11) | BioLegend | Cat# 155873; RRID: AB_2819911 |
| TotalSeq-C0308 anti-mouse Hashtag 8 (M1/42; 30-F11) | BioLegend | Cat# 155875; RRID: AB_2819912 |
| TotalSeq-C0301 anti-mouse<br>Hashtag 1 (M1/42; 30-F11) | BioLegend | Cat# 155861; RRID:<br>AB_2800693 |
| TotalSeq-C0302 anti-mouse<br>Hashtag 2 (M1/42; 30-F11) | BioLegend | Cat# 155863; RRID:<br>AB_2800694 |
| TotalSeq-C0303 anti-mouse<br>Hashtag 3 (M1/42; 30-F11) | BioLegend | Cat# 155865; RRID:<br>AB_2800695 |

**Table 6:** Anti-human antibodies used in this study.

Human
| Reagent or Resource | Source | Identifier |
| --- | --- | --- |
| Mouse anti-CD163 (clone 10D6) | Leica Biosystems | Cat# CD163-L-CE |
| Rabbit anti-VSIG4 | Novus Biologicals | Cat# NBP2-99947 |
| Donkey anti-mouse AF555 Plus | Invitrogen | Cat# A32773; RRID:<br>AB_2762848 |
| Donkey anti-rabbit AF647 Plus | Invitrogen | Cat# A32795; RRID:<br>AB_2762835 |
| Wheat Germ Agglutinin Fluorescein | Vector Laboratories | Cat# FL-1021 |

## Acknowledgements

We are very grateful for expert technical assistance of Katja Grawe, Anita Imm, Reem Alsumati, and Adriana Greco. We are indebted to the Lighthouse Core Facility at the Center for Chronic Immunodeficiency of the Medical center at the University of Freiburg, as well as the Center for Experimental Models and Transgenic Services in Freiburg for their instrumental technical assistance in flow cytometry, microscopy, and mouse work. The Lighthouse Core Facility is funded in part by the Medical Faculty, University of Freiburg (2023/A2-Fol; 2023/B3- Fol) and the Deutsche Forschungsgemeinschaft (DFG, German Research Foundation) (450392965).

Additionally, we like to thank the BioMaterialBank Nord for providing human samples.

The BioMaterialBank Nord is supported by the German Center for Lung Research. The BioMaterialBank Nord is member of popgen 2.0 network (P2N).

FL and AKL were recipients of the Clinician scientist program IMM-PACT stipend, funded by the DFG (413517907).

CS has received funding through the SFB1453 (431984000), SCHE 2092/4-1 (RP9, CP2, CP3) (241702976 and 438496892), SFB1160 (project-ID 256073931), and the Heisenberg program (501370692).

Funding to SP was provided by the DFG (491676693 TRR 359 - PILOT).

Sagar is supported by the Department of Medicine II, Freiburg University Medical Center, Faculty of Medicine, University of Freiburg and by the DFG (491676693 TRR 359 - PILOT).

PH has received funding by the DFG (283781347, 254895677, 446317895-SPP EXIT, 259373024 - TRR 167 NeuroMac, and 491676693 TRR 359 - PILOT).

## Contributions

PH and JN conceptualized the study and wrote the manuscript. JN performed most of the experiments and prepared the figures. FL helped with the analysis and provided critical intellectual input.

SW performed the ATAC sequencing under supervision of SP and helped with the analysis. MG helped with the establishment of the Kupffer cell culture.

AKL provided critical intellectual input.

DO helped with the trajectory analysis under the supervision of S. S performed the scRNA sequencing and helped with its analysis.

VG performed retro-orbital infections.

TG provided human samples of patients infected with *Mtb* and performed the H&E staining. MR and CS established the immunofluorescence staining of the human livers and CS confirmed granuloma presence.

All co-authors have read and edited the manuscript.

