## Supplementary Figures for "Mycobacterial infection uncovers plasticity of Kupffer cells"

Extended Data Figure 1

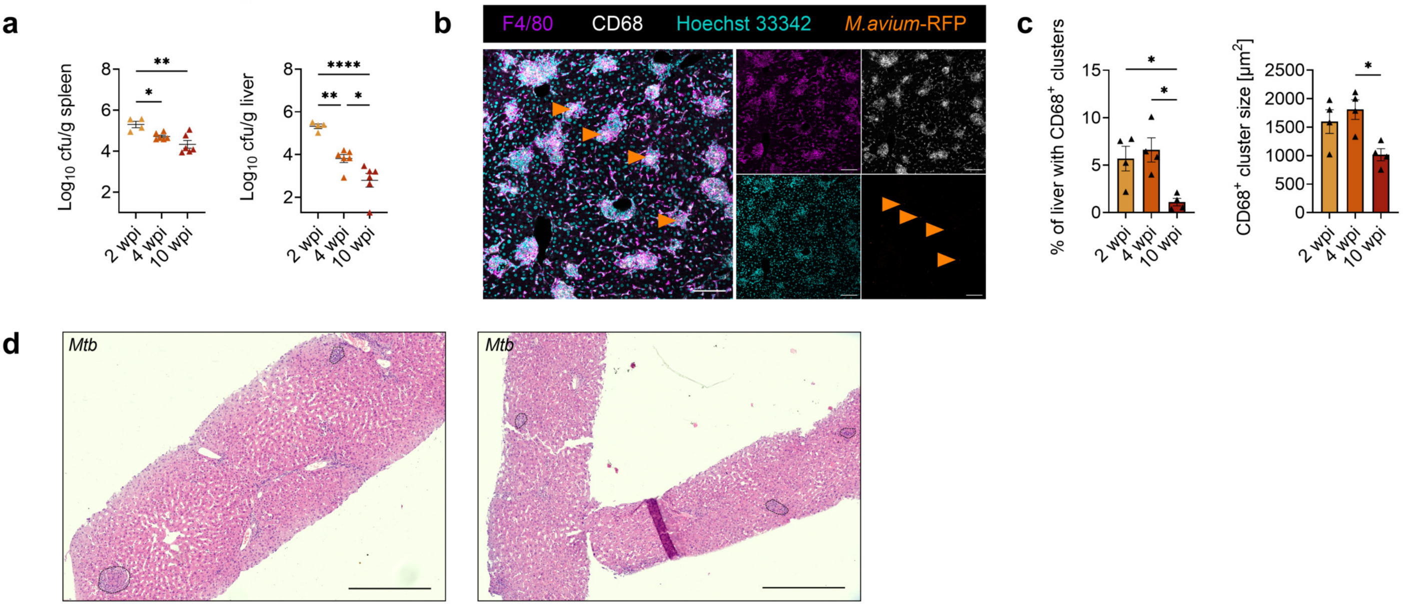

**Extended Data Figure 1: Granuloma formation in mycobacterial infections**

(a) Bacterial burden in spleen and liver at indicated time points after *M. avium*-RFP infection of C57BL/6J mice. n=(4, 6, 6) mice respectively. One mouse at 10 wpi was below the detection limit in the liver.

(b) Immunofluorescence staining of a C57BL/6J mouse liver 2 weeks after *M. avium*-RFP infection. Orange arrowheads indicate *M. avium* within granulomas. Scale bars: 100 µm.

(c) Quantification for the liver area occupied by granulomas, defined as $CD68^+$ clusters with a minimal size of 350 $\mu m^2$, as well as their average size based on the microscopy images shown in (b). n=4 per group.

(d) Liver overviews of an H&E staining from two patients with confirmed *Mtb* infection (related to Figure 1I). Black dashed lines indicate granulomas. Scale bar: 500 µm.

Data represent mean ± SEM with each symbol depicting one mouse are derived from at least 2 independent experiments. One-way ANOVA with Tukey‘s multiple comparisons test. * $P \le 0.05$, ** $P \le 0.01$, *** $P \le 0.001$, **** $P \le 0.0001$.

Extended Data Figure 2

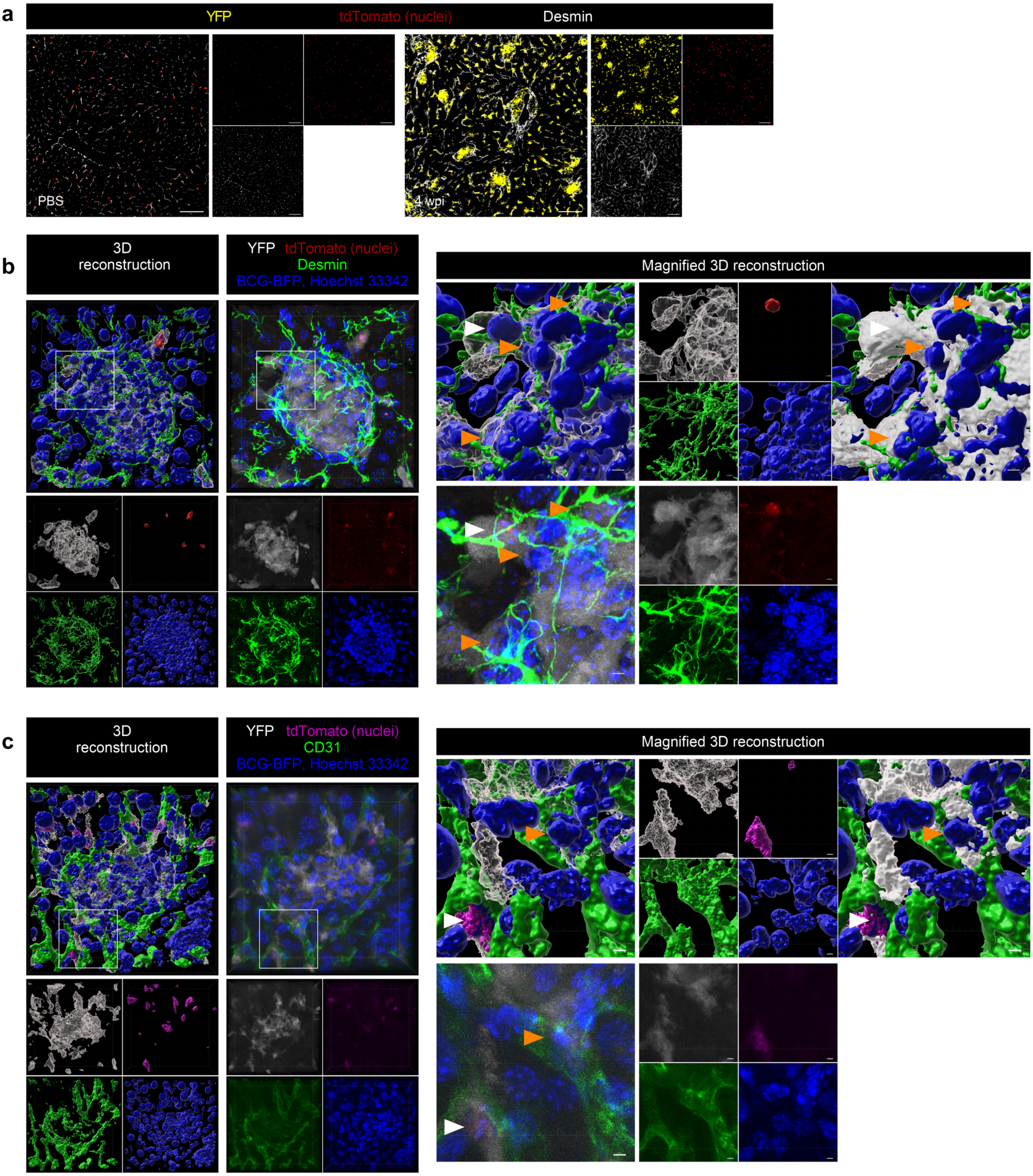

**Extended Data Figure 2: Interactions of $KC^{low}$ with hepatic stellate cells and liver sinusoidal endothelial cells**

(a) Immunofluorescence (IF) staining of *Clec4f*$^{Cre\text{-}tdTomato\text{-}NLS}$*:ROSA26*$^{EYFP}$ livers of PBS injected mice and at 4 wpi with BCG-BFP. Depicted are the reporters YFP (yellow), and tdTomato (red), as well as a staining against desmin (white) to label hepatic stellate cells. Scale bars: 100 µm.

(b) 3D reconstruction and IF of a granuloma from a *Clec4f*$^{Cre\text{-}tdTomato\text{-}NLS}$*:ROSA26*$^{EYFP}$ mouse 4 weeks after infection with BCG-BFP stained against desmin (green) and Hoechst 33342 (blue). YFP is depicted in white, while tdTomato is in red. Images on the right display a magnified view of the indicated area on the left. Arrowheads mark close cell contacts of $KC^{high}$ (white arrowheads) or $KC^{low}$ (orange arrowheads) with hepatic stellate cells within the granuloma. Scale bars: Overview 10 µm per major tick, magnified picture 3 µm.

(c) 3D reconstruction and IF of a granuloma from a *Clec4f*$^{Cre\text{-}tdTomato\text{-}NLS}$*:ROSA26*$^{EYFP}$ mouse 4 weeks after infection with BCG-BFP stained against CD31 (green) and Hoechst 33342 (blue). CD31 was used to label liver sinusoidal endothelial cells. YFP is depicted in white, while tdTomato is in magenta. Images on the right display a magnified view of the indicated area on the left. Arrowheads mark close cell contacts of $KC^{high}$ (white arrowheads) or $KC^{low}$ (orange arrowheads) with liver sinusoidal endothelial cells within the granuloma. Scale bars: Overview 10 µm per major tick, magnified picture 3 µm.

Extended Data Figure 3

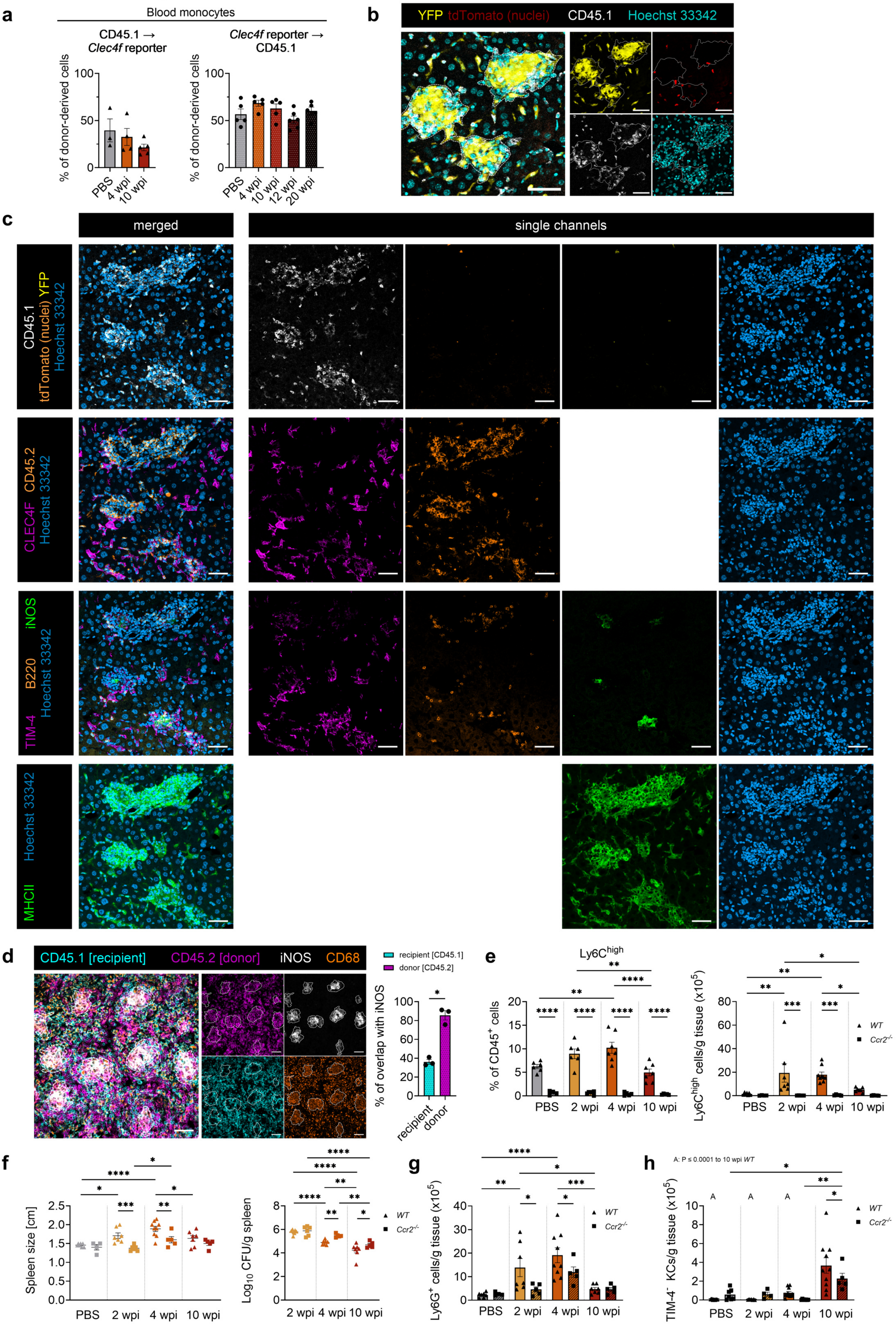

**Extended Data Figure 3: Contribution of bone marrow-derived monocytes to granuloma formation**

(a) Blood chimerism of donor-derived monocytes in *Clec4f*$^{Cre\text{-}tdTomato\text{-}NLS}$*:ROSA26*$^{EYFP}$ mice transplanted with CD45.1 bone marrow or vice versa. Mice are derived from Fig. 2b.

(b) Liver immunofluorescence staining (IF) of a *Clec4f*$^{Cre\text{-}tdTomato\text{-}NLS}$*:ROSA26*$^{EYFP}$ mouse transplanted with CD45.1 bone marrow at 4 wpi. Dotted line indicates granulomas. Scale bar: 50 µm

(c) Liver multicycle IF staining from a CD45.1 mouse transplanted with *Clec4f*$^{Cre\text{-}tdTomato\text{-}NLS}$*:ROSA26*$^{EYFP}$ bone marrow at 4 wpi. Each staining round is represented in a separate row. YFP and tdTomato (first row) were stained over in subsequent rounds. Scale bars: 50 µm.

(d) IF of a spleen from a CD45.1 mouse transplanted with *Clec4f*$^{Cre\text{-}tdTomato\text{-}NLS}$*:ROSA26*$^{EYFP}$ bone marrow at 4 wpi and the quantification for the overlap of CD45.1 or CD45.2 with iNOS based on microscopic images. n=3 per group. Dotted lines mark granulomas. Scale bar: 50 µm.

(e) Flow cytometry analysis of Ly6C$^{high}$ blood (left) and liver (right) monocytes in *WT* ($n_{blood}$=(6, 6, 7, 7), $n_{liver}$=(7, 7, 9, 7) in sequence) and *Ccr2*$^{-/-}$ ($n_{blood}$=(4, 6, 5, 5), $n_{liver}$=(4, 5, 5, 5) in sequence) mice in infection.

(f) Spleen size and bacterial burden comparing *WT* (n=(7, 7, 9, 7) in sequence) and *Ccr2*$^{-/-}$ (n=(4, 6, 5, 5) in sequence) mice after infection.

(g) Flow cytometry of liver granulocytes comparing *WT* and *Ccr2*$^{-/-}$ mice from (e).

(h) Flow cytometry of TIM-4$^{-}$ KCs in *Clec4f*$^{Cre\text{-}tdTomato\text{-}NLS}$*:ROSA26*$^{EYFP}$ and *Clec4f*$^{Cre\text{-}tdTomato\text{-}NLS}$*:ROSA26*$^{EYFP}$*:Ccr2*$^{-/-}$ mice. Mice are derived from Fig. 2j.

Experiments (a-g) were performed with WT BCG. Data represent mean ± SEM with each symbol depicting one mouse and are derived from at least 2 independent experiments. One-way ANOVA (a), two-tailed paired t test (d), or two-way ANOVA (e-h) with Tukey‘s multiple comparisons test. * $P \leq 0.05$, ** $P \leq 0.01$, *** $P \leq 0.001$, **** $P \leq 0.0001$.

### Extended Data Figure 4

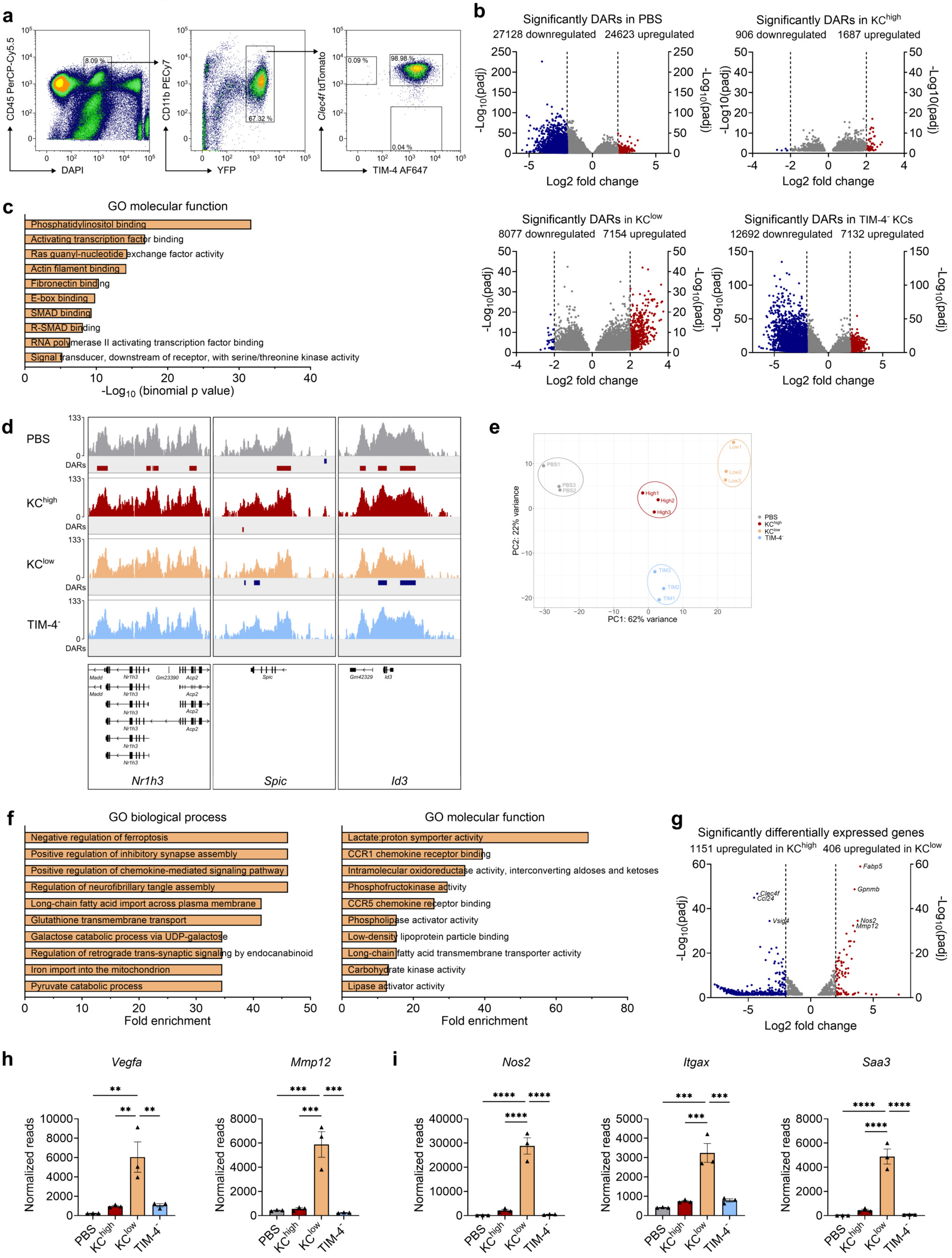

**Extended Data Figure 4: Epigenetic and transcriptional adaptations of KCs**

(a) Representative sorting strategy for ATACseq samples depicted for a PBS treated mouse. Sorted populations were KC$^{high}$, KC$^{low}$, and TIM-4$^{-}$ KCs in BCG-BFP infected or PBS treated *Clec4f*$^{Cre\text{-}tdTomato\text{-}NLS}$:*ROSA26*$^{EYFP}$ mice. Percentages of the gated populations are depicted in the graph. Related to Figure 3a.

(b) Volcano plot of significantly differential accessible regions (DARs) for each population compared to all others in the ATACseq data. Coloured genes are above the $\log_2$ fold change threshold of 2.

(c) Top 10 enriched GO-terms of molecular functions for KC$^{low}$ vs. all other populations in the ATACseq data.

(d) Integrative Genomics Viewer of ATACseq displaying chromatin accessibility for indicated genes representative for one sample per population. Significantly DARs are depicted below the tracks in blue for downregulated and red for upregulated accessibility comparing one population vs. all other populations. Tracks for *Madd* were simplified.

(e) PCA plot from bulk RNAseq of sorted KC$^{high}$, KC$^{low}$, and TIM-4$^{-}$ KCs of *Clec4f*$^{Cre\text{-}tdTomato\text{-}NLS}$:*ROSA26*$^{EYFP}$ mice 4 weeks after BCG-BFP infection, as well as KCs from PBS treated mice. n = 3 per group.

(f) Top 10 GO terms for differentially expressed genes (DEGs) upregulated in KC$^{low}$ compared to KC$^{high}$ using the bulk RNAseq data.

(g) Volcano plot of significantly DEGs comparing KC$^{high}$ to KC$^{low}$ in bulk RNAseq. Coloured genes are above the $\log_2$ fold change threshold of 2.

(h, i) Normalized reads for indicated genes in KC subsets in the bulk RNAseq data. Data represent mean ± SEM with each symbol depicting one mouse. One-way ANOVA with Tukey‘s multiple comparisons test. * $P \leq 0.05$, ** $P \leq 0.01$, *** $P \leq 0.001$, **** $P \leq 0.0001$.

Extended Data Figure 5

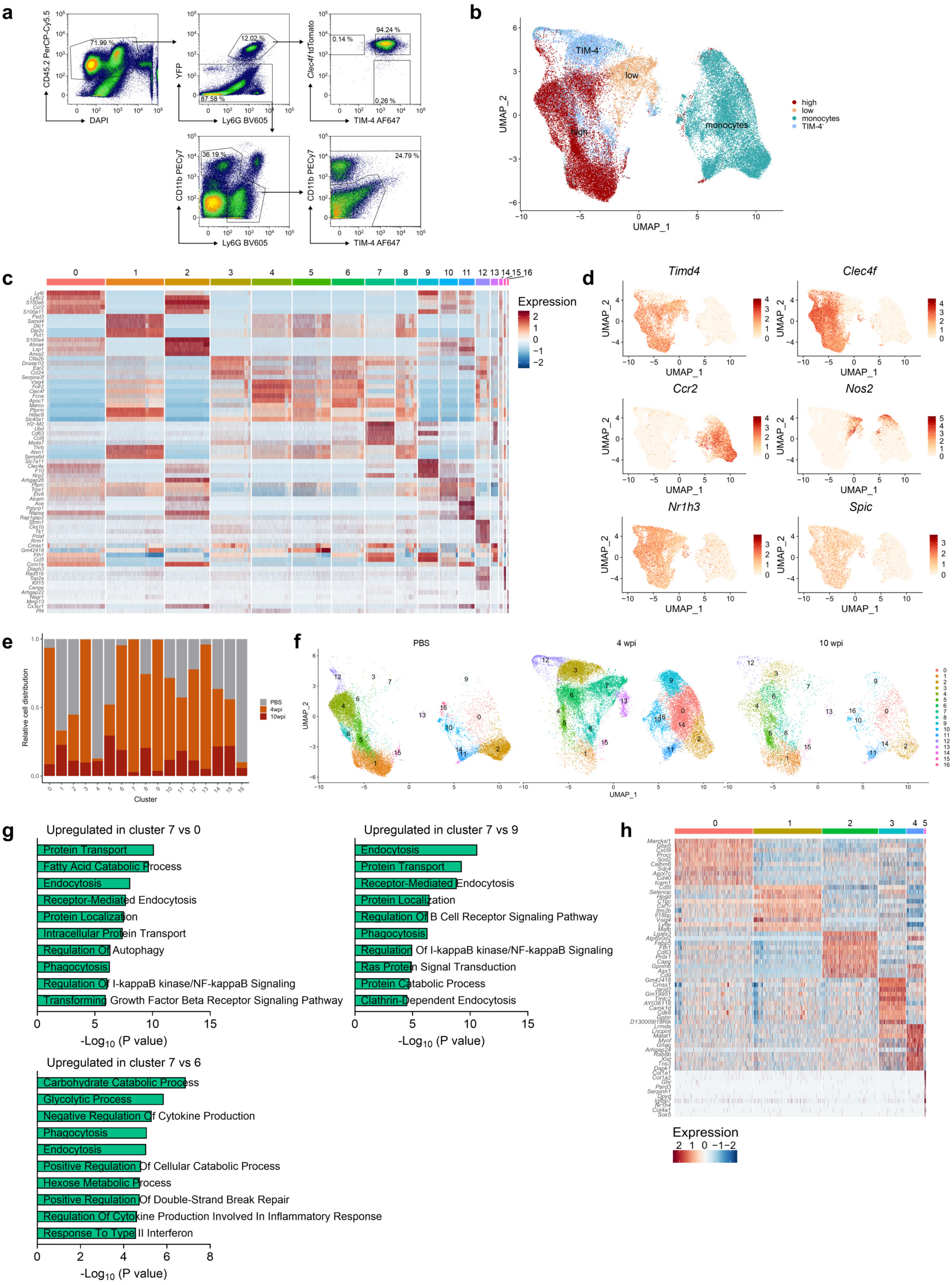

**Extended Data Figure 5: Identification of infection specific KC clusters**

(a) Sorting strategy for the scRNA sequencing experiment, exemplary shown for a PBS injected mouse. $KC^{high}$, $KC^{low}$, TIM-$4^{-}$ KCs, and $CD11b^{high}$ or TIM-$4^{+}$ cells were sorted separately from *Clec4f*$^{\textit{Cre-tdTomato-NLS}}$*:ROSA26*$^{\textit{EYFP}}$ mice after PBS injection or 4 and 10 weeks after BCG-BFP infection. Percentages of the gated populations are depicted in the graph.

(b) UMAP with colour coding based on the sorted population after removal of contaminating cells. Monocytes were sorted as the $CD11b^{high}$ or TIM-$4^{+}$ cell population. In total 48,500 cells.

(c) Heatmap with the top 5 differentially expressed genes calculated for each cluster.

(d) Feature plot showing the expression of KC signature genes, *CCR2* and *Nos2*.

(e) Relative cell distribution per time point (PBS, 4 wpi, or 10 wpi) of the identified clusters.

(f) UMAP split by the time points (PBS, 4 wpi, and 10 wpi).

(g) GO terms for biological processes upregulated in cluster 7 ($KC^{low}$) compared to cluster 0, 6, or 9.

(h) Heatmap for the subclustered $KC^{low}$ dataset (Figure 4i) with the top 10 differentially expressed genes calculated for each cluster. In total 3183 cells.

Extended Data Figure 6

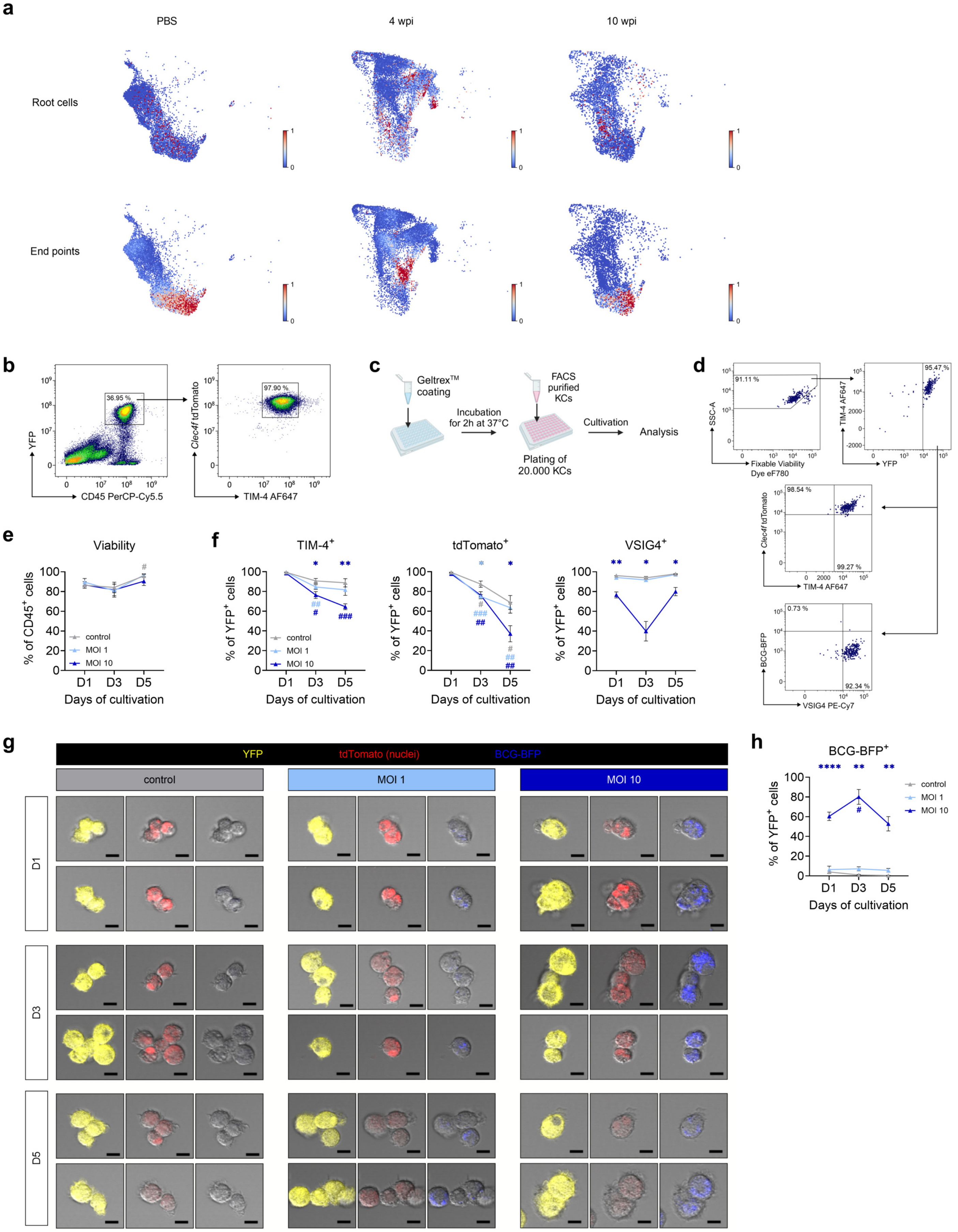

**Extended Data Figure 6: Cultivated KCs are viable and maintain their markers**

(a) Root cells and end points determined within the trajectory analysis in Figure 5a split by condition.

(b) Sorting strategy of KCs from $\textit{Clec4f}^{\textit{Cre-tdTomato-NLS}}\textit{:ROSA26}^{\textit{EYFP}}$ mice for *in vitro* cultivation, pre-gated on single cells.

(c) Experimental setup for *in vitro* cultivation of sorted KCs from (a), plated on Geltrex™ coated plates. Cultured KCs were treated with BCG-BFP at MOI 1, MOI 10, or left untreated.

(d) Gating strategy used in (d, e, g), shown for the control condition on day 1 after cultivation, pre-gated on live $CD45^+$ single cells.

(e) KC viability during cultivation assessed by flow cytometry. n = 5 per group.

(f) Flow cytometry of TIM-4, tdTomato, and VSIG4 expression in KCs from (d).

(g) Confocal microscopy of cultured KCs with two representative pictures (top and bottom) per condition and cultivation day. Bright fields are overlayed with YFP (yellow, left), tdTomato (red, middle), and BCG-BFP (blue, right). Scale bar: 10 µm.

(h) Flow cytometry of $BFP^+$ KCs due to phagocytosed BCG, related to (d).

Data represent mean ± SEM and are derived in (d-h) from 2 independent experiments. Percentages of the gated populations are depicted in the graph. Statistical significance was calculated in (c-e) between the condition and control at the same day (asterisks), and during the cultivation for a single condition compared to D1 (hashtags) using a two-way ANOVA analysis with Tukey's multiple comparison test. * $P \leq 0.05$, ** $P \leq 0.01$, *** $P \leq 0.001$, **** $P \leq 0.0001$. D, day.

### Supplemental Figure 1

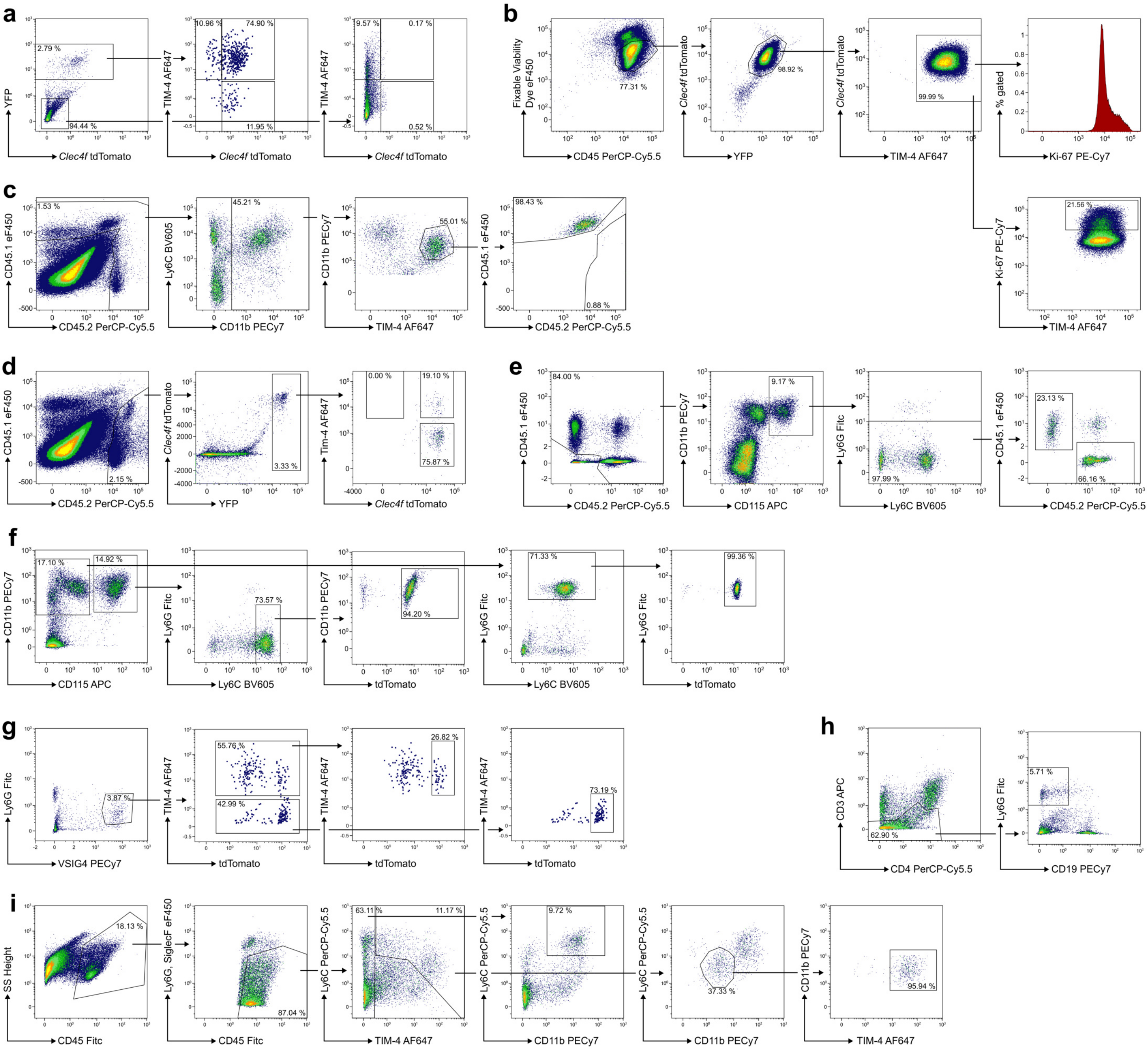

**Supplemental Figure 1: Exemplary flow cytometry gating strategies**

(a) KC subsets in a $Clec4f^{Cre\text{-}tdTomato\text{-}NLS}$:$ROSA26^{EYFP}$ mouse 4 wpi with BCG-BFP. YFP$^{-}$ cells were gated as a negative control to adjust the gate for the tdTomato expression. Related to Figure 1e, f, h.

(b) Ki-67 staining of sorted KC$^{high}$ from a $Clec4f^{Cre\text{-}tdTomato\text{-}NLS}$:$ROSA26^{EYFP}$ mouse 2 wpi with BCG-BFP. Related to Figure 1g.

(c) Gating of TIM-4$^{+}$ KCs shown for a PBS injected CD45.1 mouse transplanted with $Clec4f^{Cre\text{-}tdTomato\text{-}NLS}$:$ROSA26^{EYFP}$ bone marrow. Related to Figure 2b.

(d) Gating of KC subsets shown for a CD45.1 mouse transplanted with $Clec4f^{Cre\text{-}tdTomato\text{-}NLS}$:$ROSA26^{EYFP}$ bone marrow 10 wpi with BCG WT. Related to Figure 2e.

(e) Gating of blood monocytes shown for a PBS injected CD45.1 mouse transplanted with $Clec4f^{Cre\text{-}tdTomato\text{-}NLS}$:$ROSA26^{EYFP}$ bone marrow. Related to Extended Data Figure 3a.

(f) Gating of tdTomato expression in blood for Ly6C$^{high}$ monocytes and Ly6G$^{+}$ granulocytes for an $Ms4a3^{Cre}$:$ROSA26^{tdTomato}$ mouse 4 wpi with BCG-BFP. Related to Figure 2f. A similar gating was used for Extended Data Figure 3e.

(g) Gating of tdTomato expression in TIM-4$^{+}$ and TIM-4$^{-}$ KCs of an $Ms4a3^{Cre}$:$ROSA26^{tdTomato}$ mouse 10 wpi with BCG-BFP. Related to Figure 2g.

(h) Granulocyte gating in a *WT* mouse 4 wpi with BCG WT. Related to Extended Data Figure 3g.

(i) Ly6C$^{high}$ monocyte and TIM-4$^{+}$ KCs gating in a *WT* mouse 4 wpi with BCG WT. Related to Extended Data Figure 3e, h.

Cells were pre-gated as singlets (b-e, i) and CD45$^{+}$ cells (a, f-h). Unless otherwise specified livers were used for analysis. Percentages of the gated populations are depicted in the graphs.
